# An Advanced Humanized Systemic Lupus Erythematosus Model Enables Parallel Profiling of B Cell-Targeted Therapies

**DOI:** 10.64898/2026.01.27.701893

**Authors:** Rujie Zhu, Shuai Ding, Deshan Ren, Juan Liang, Shuang Liu, Jialu Fan, Shuhua Yu, Zijian Zhang, Chunyu Cheng, Kai Wang, Yuxuan Chen, Yang Hang, Xiaoyun Shang, Yan Li, Lingyun Sun

**Affiliations:** Department of Rheumatology and Immunology, Nanjing Drum Tower Hospital, Nanjing Drum Tower Hospital Clinical College of Nanjing Medical University, Nanjing, 211166, China; State Key Laboratory of Reproductive Medicine and Offspring Health, Nanjing Medical University, Nanjing, 211166, China; MOE Key Laboratory of Model Animal for Disease Study, Model Animal Research Center, Department of Rheumatology and Immunology, Nanjing Drum Tower Hospital, Affiliated Hospital of Medical School, Nanjing University, Nanjing 210061, China; Department of Rheumatology and Immunology, Nanjing Drum Tower Hospital, The Affiliated Hospital of Nanjing University Medical School, Nanjing University, Nanjing, 211166, China; GemPharmatech Co., Ltd., 12 Xuefu Road, Jiangbei New Area, Nanjing, 210032, China; Department of Rheumatology and Immunology, The Affiliated Huaian No.1 People’s Hospital of Nanjing Medical University, Huaian, 223001, China; T-Maximum Pharmaceutical (Suzhou) Co., Ltd., Suzhou, 215028, China; Chongqing International Institute for Immunology, Chongqing, 401338, China; Changping Laboratory, Beijing, 102200, China; ChemBioMed Interdisciplinary Research Center at Nanjing University, Nanjing, 210061, China; Wuxi Xishan NJU Institute of Applied Biotechnology, Wuxi, 214101, China

**Keywords:** systemic lupus erythematosus, humanized mouse model, B cell-targeted therapy, CAR-T cells, T-cell engagers, translational immunology

## Abstract

B cell-targeted therapies represent a transformative frontier for systemic lupus erythematosus (SLE) intervention, yet clinical recommendation of specific treatment is hampered by the challenge to perform head-to-head comparisons of efficacy-toxicity trade-offs and tissue-specific impacts in patients. To address this gap, an advanced Toll-like receptor 7 (TLR7) agonist-induced SLE model is developed in a NCG-M (NOD-*Prkdc^em26Cd52^Il2rg^em26Cd22^Rosa26^em1Cin(hCSF2&IL3&KITLG)^*/Gpt) human immune system (HIS) mice. This model enables parallel evaluation of multiple B cell-directed therapies within a controlled cohort study. It recapitulates core SLE pathologies—including immune effector cell augmentation, autoantibody production and glomerulonephritis—within 2 months, demonstrating greater severity and clinical fidelity than conventional pristane-induced systems. Validated through replication of clinical responses to rituximab and belimumab, the platform directly compares two emerging modalities: universal chimeric antigen receptor-T cells (UCAR-T) elicits delayed but profound B-cell reset correlating with superior reduction in renal pathology and autoantibodies, while T cell engagers (TCEs) mediate rapid yet partial B-cell depletion and transient efficacy. UCAR-T concurrently induces greater acute inflammatory responses, aligning with its distinct toxicity profile. By resolving therapy-specific effects on human immune dynamics and symptom control, this study empowers context-specific therapeutic selection for SLE, advancing tailored management strategies.

## Introduction

Systemic lupus erythematosus (SLE) is a complex autoimmune disease characterized by autoreactive immune cells, autoantibodies and subsequent multi-organ tissue damage.^1^ Although standard treatment regimens combining corticosteroids with immunosuppressants are effective for most patients, a subset with refractory, relapsing disease remains challenging to manage.^2^ Extensive evidence implicates aberrant B cell responses as central drivers of SLE pathogenesis,^3^ establishing them as key therapeutic targets. While two B cell-targeted monoclonal antibodies — rituximab (anti-CD20) and belimumab (anti-B-cell-activating factor, BAFF)—are approved for SLE treatment, both demonstrate clinical limitations. Rituximab failed primary endpoints in pivotal randomized controlled trials despite initial promise,^4–6^ while belimumab exhibits delayed efficacy in certain patient subsets.^7^

The pursuit of durable drug-free remission in SLE has accelerated development of advanced B cell-targeted therapeutics, including chimeric antigen receptor (CAR) T cells and T cell engagers (TCEs).^8^ Autologous CAR-T therapy demonstrates promising efficacy in pilot clinical trials,^9^ yet remains constrained by complex manufacturing and prohibitive financial burden. Novel strategies, including allogeneic “off-the-shelf” CAR-T products,^10^ regulatory CAR-T cell designs,^11^ and bispecific TCEs,^12^ are currently under investigation as promising alternative therapies. A pioneering study comparing clinical trials of different B cell-depleting strategies revealed significant variations in depletion efficiency between protein-based and cell-based therapies.^13^ While this investigation represents the first reported comparison of multiple B cell-targeted therapeutics in autoimmune diseases, its clinical implications were constrained by several limitations: patient heterogeneity, restricted tissue accessibility, and a retrospective design that pooled data from trials on different diseases and cohorts. As such, a controlled cohort study directly evaluating efficacy, toxicity, and multi-tissue-level pharmacodynamics in SLE would provide valuable insight and evidence for selecting between these emerging treatment options.

Human immune system (HIS) mice offer a valuable platform to overcome these challenges.^14^ Patient peripheral blood mononuclear cell (PBMC)-humanized lupus models were constrained by poor B cell reconstitution, limited cell diversity, and rapid onset of graft-versus-host disease (GVHD), restricting their utility for evaluating B cell-targeted therapies.^15,16^ In contrast, hematopoietic stem cell (HSC)-humanized mice support broader human immune cell development and long-term studies.^17^ The initial HSC-humanized lupus model, generated through intraperitoneal injection of pristane into NSG (NOD/LtSz-*Prkdc^scid^Il2rg^tm1whl^*/J) HIS mice,^18^ induced peritoneal inflammation and demonstrated lupus-like phenotypes. Although representing a significant advancement, insufficient activation of the type I interferon (IFN-I) signaling pathway and attenuated autoantibody responses limit its effectiveness for evaluating B cell-targeted therapeutic interventions.

Given these challenges, we sought to establish a humanized lupus model centered on the TLR7-IFN-I signaling axis, a key driver of SLE pathogenesis. Gain-of-function mutations in TLR7 have been identified as a monogenic cause of lupus,^19^ and clinical reports have documented drug-induced SLE in women receiving low-dose topical TLR7 agonists for unrelated indications.^20^ To recapitulate this pathway *in vivo*, we employed a topical TLR7 agonist to achieve titratable and reproducible activation of IFN-I signaling. In parallel, we screened and selected a next-generation immunodeficient mouse strain (NCG-M, NOD-*Prkdc^em26Cd52^Il2rg^em26Cd22^Rosa26^em1Cin(hCSF2&IL3&KITLG)^*/Gpt) engineered to express three human cytokines that support robust human myeloid lineage development, B cell maturation, and immunoglobulin production after human HSC engraftment. This dual-optimization approach — targeted TLR7 activation and host strain enhancement — provides a biologically relevant foundation for modeling human SLE. Building on this system, we designed a head-to-head preclinical evaluation comparing two approved monoclonal antibodies and two emerging B cell-redirected immunotherapies—universal CAR-T (UCAR-T) cells and TCEs, aiming to guide therapy optimization and precision medicine development in SLE.

## Results

### Topical TLR7 agonist treatment triggers lupus-like phenotype in NCG HIS mice

Previous studies have shown that epicutaneous application of TLR7 agonists induces systemic autoimmunity in wild-type FVB/N, BALB/c, and C57BL/6 mice.^21,22^ To assess whether epicutaneous TLR7 agonists induce systemic autoimmunity in a human immune context similarly to conventional mouse strains, we engrafted human HSCs into immunodeficient NCG (NOD-*Prkdc^em26Cd52^Il2rg^em26Cd22^*/Gpt) mice to generate NCG HIS mice. Newborn immunodeficient NCG mice (4-6 days old) were irradiated with X-ray (80cGy) and injected intra-hepatically with 1 × 10^5^ human HSCs.^23^ Successful engraftment was confirmed by peripheral blood analysis, with human white blood cells (WBCs) exceeding 2 × 10^5^/mL and accounting for more than 20% of total WBCs 12 weeks post-engraftment (Figure S1a&1b).

Qualifying NCG HIS mice meeting these engraftment criteria subsequently received the initial induction protocol: thrice-weekly epicutaneous administration of 100μg Resiquimod (R848), a TLR7 agonist, applied near the marginal ear vein (Figure S1c). Within two weeks, peripheral blood showed a dramatic reduction in human immune cells (Figure S1d). Flow cytometry revealed transitional B cells exceeding 90% of total B cells in peripheral blood from TLR7 agonist-induced mice, with comparable distributions in spleen and bone marrow as shown in Figure S1e. These findings suggest marked activation-induced cell death and compensatory immature B cell mobilization from bone marrow, indicating that human immune cells are hyperresponsive to TLR7 agonists *in vivo*. To mitigate this effect, we reduced the dose of TLR7 agonist to one fifth of initial treatment (Figure 1A) and monitored the mice every two weeks for two months. Under this regimen, human immune cell counts and percentages remained stable with minimal decreases (Figure S2a). Following two months of induction, erythema appeared on the faces of the model group, recapitulating a characteristic feature of SLE patients (Figure 1B). Subsequent immune cell phenotyping confirmed immune dysregulation consistent with lupus pathogenesis. In TLR7 agonist-induced group, T cells exhibited increased activation (by HLA-DR) and a shift from naïve to effector phenotypes (via CD45RA and CCR7 patterns) (Figure 1C, Figure S2b&2c), mirroring T cell profiles observed in SLE patients (Figure 1D). Treated HIS mice exhibited a pronounced Th1/Th17 polarization, demonstrating significantly elevated production of pro-inflammatory cytokines (IFN-γ, IL-17A, and IL-21) upon stimulation (Figure 1E, Figure S2d), consistent with clinical observations in SLE patients (Figure 1F). Furthermore, TLR7 agonist-challenged HIS mice recapitulated the characteristic cytokine profile of human SLE (Figure 1H), showing marked increases in serum IL-6 and IL-10 levels that closely paralleled those detected in patient samples (Figure 1G).

**Figure 1.**
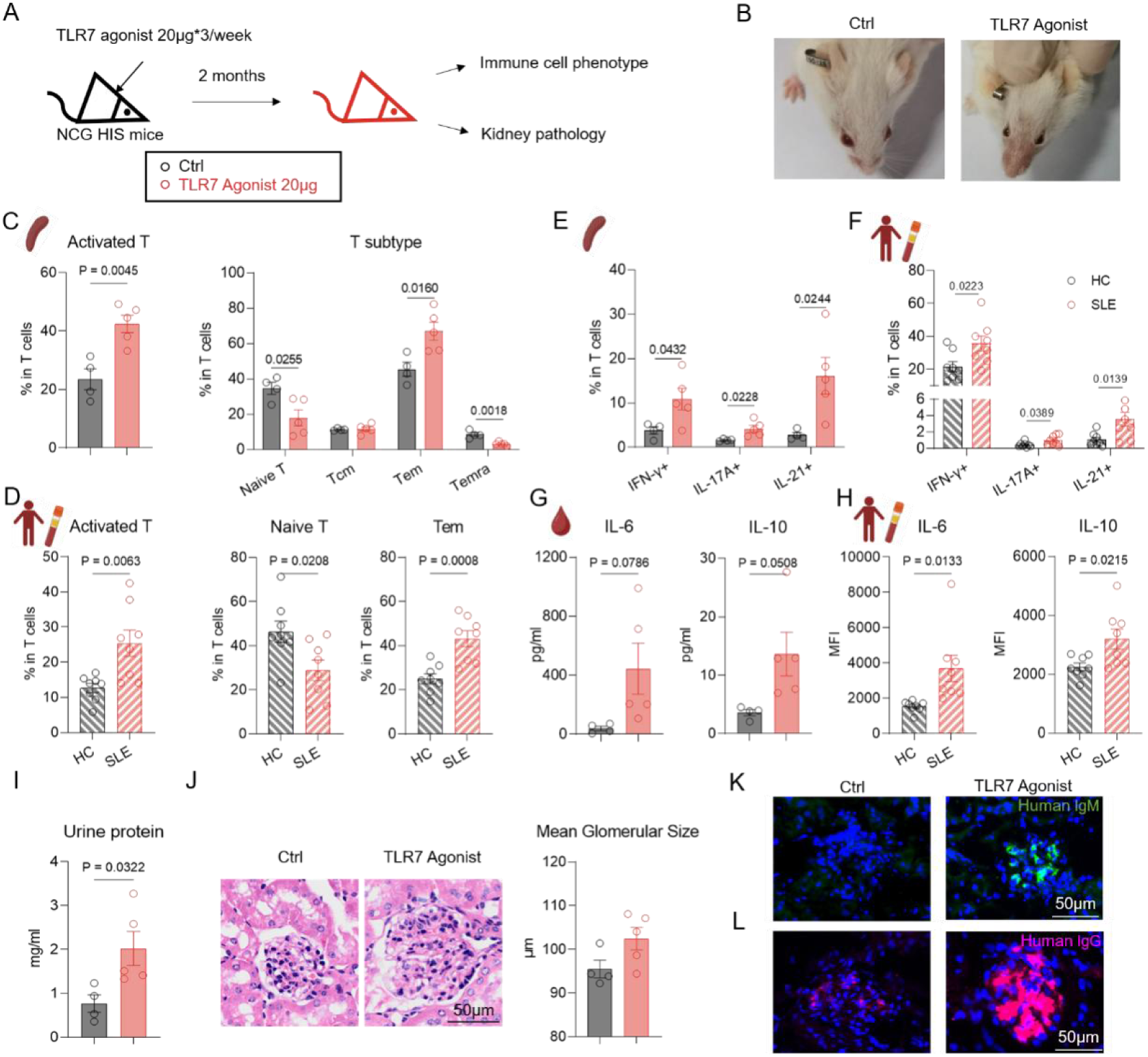
Topical TLR7 agonist induction partially recapitulates SLE clinical pathology in NCG HIS mice. A) NCG HIS mice were topically treated with low dose TLR7 agonist (20μg*3/week) for 8 weeks, disease phenotypes were determined at the endpoint. Ctrl group, *n=4*. TLR7 agonist (20μg)-induced group, *n=5*. B) Representative images of facial erythema in two groups. C-D) Phenotypic analysis of T cells in spleens from NCG HIS mice (C) and peripheral blood from human (D). Frequencies of HLA-DR^+^ (activated), CD45RA^+^CCR7^+^ (naive), CD45RA^+^CCR7^−^(Temra), CD45RA^−^CCR7^+^ (Tcm) and CD45RA^−^CCR7^−^ (Tem) cells were summarized. E-F) Phenotypic analysis of T cell subsets in spleens from NCG HIS mice (E) and peripheral blood from human (F). Frequencies of IFN-γ^+^, IL-17A^+^ and IL-21^+^ cells were summarized. G-H) Serum cytokine levels from NCG HIS mice(G) and human(H). I-J) Kidney pathology in NCG HIS mice. I) Protein levels in urine from control and TLR7 agonist-induced mice. J) Representative HE staining images of glomerular area in the kidney tissue. Mean size of glomerular was summarized. Scare bar, 50μm. K-L) Immunofluorescence staining of human IgM (K) and human IgG (L) deposition in the glomerular area of kidneys from control and TLR7 agonist-induced mice. Scare bar, 50μm. Data show mean ± SEM. Statistical analysis was performed using unpaired two-tailed *t*-test. The exact *p* values are shown.

Chronic immune imbalance ultimately led to tissue pathology. The most prominent manifestation was lupus nephritis, characterized by multiple immune complex deposition within glomeruli, provoking local inflammation and consequent tissue damage.^24^ TLR7 agonist-induced NCG HIS mice also exhibited excess urinary protein (Figure 1I), glomerular endothelial and mesangial hypercellularity (Figure 1J), and IgG and IgM deposition (Figure 1K&1L), recapitulating human lupus nephritis. While our model successfully captured T cell dysregulation and renal pathology through optimized TLR7 agonist dosing, it exhibited limitations in humoral immunity. Conventional HIS mice showed impaired B cell function and low baseline serum IgG levels,^25^ resulting in undetectable autoantibodies despite a >10-fold induction of serum IgG after R848 treatment (absolute concentration: 10μg/mL; 0.1% of normal human levels; Figure 2A).

**Figure 2:**
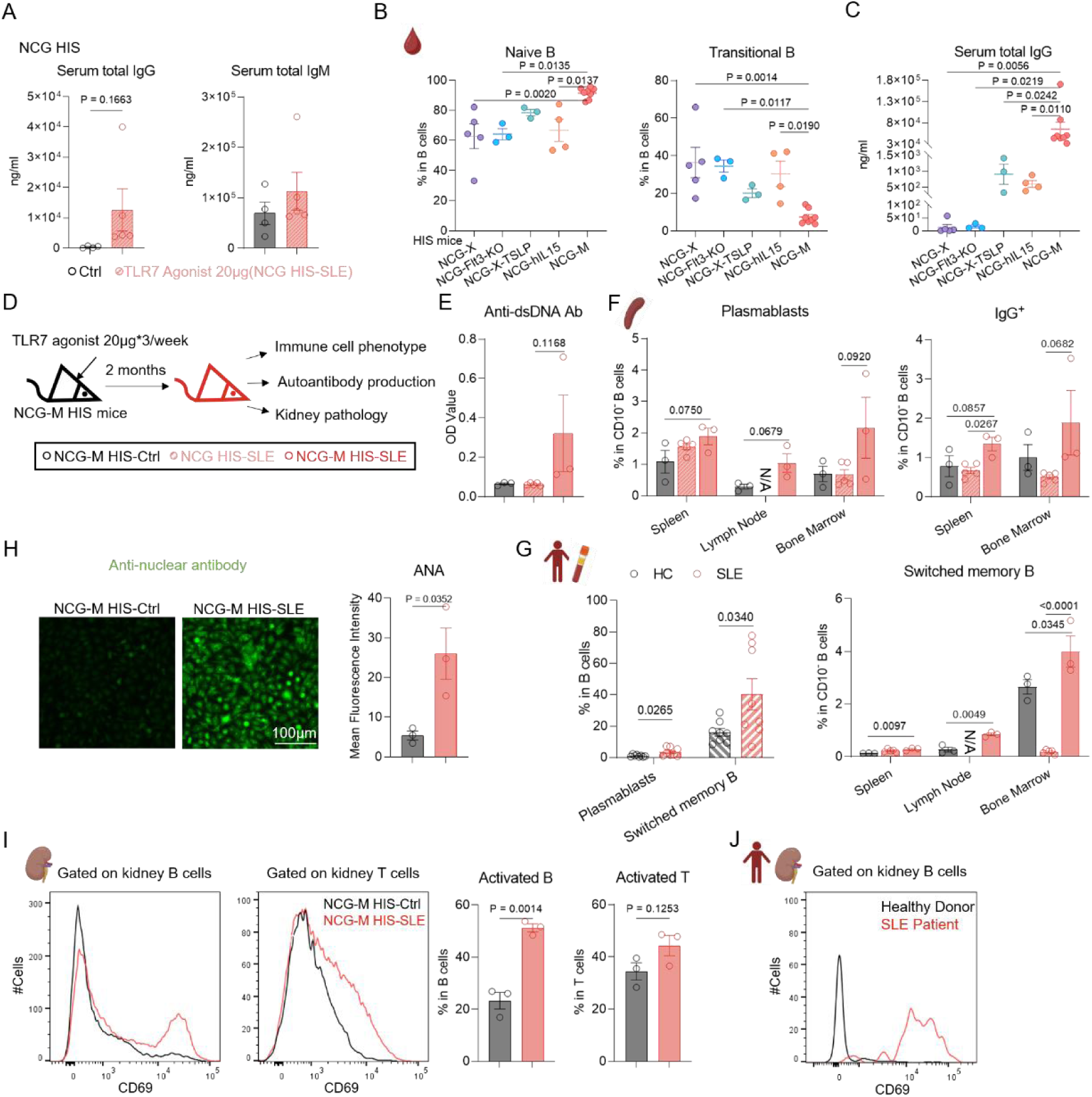
Lupus-like pathology mediated by autoreactive B cells in NCG-M HIS mice. A) Total IgG and IgM level in serum from Ctrl*(n=4)* and TLR7 Agonist-induced *(n=5)* group (NCG HIS mice) detected by ELISA. B) Flowcytometry analysis of B cell subsets in peripheral blood from different HIS mice background. C) Total IgG level in serum from different HIS mice background detected by ELISA. D) NCG-M HIS mice were topically induced with low dose TLR7 agonist (20μg*3/week) for 8 weeks, disease phenotypes were determined at the endpoint. NCG-M HIS-Ctrl group, *n=3*. NCG-M HIS-SLE group, *n=3*. NCG HIS-SLE group, *n=5*. E) Serum anti-dsDNA antibody detected by ELISA. F-G) Phenotypic analysis of B cells in three tissues from NCG HIS or NCG-M HIS mice (F) and peripheral blood from human (G). Frequencies of CD24^−^CD38^hi^ (plasmablast), CD27^+^IgD^−^ (switched memory, SWM) and CD27^+^ IgG^+^ cells were summarized. H) Representative images and summarized data of serum anti-nuclear antibody in NCG-M HIS-Ctrl and NCG-M HIS-SLE group detected by indirect immunofluorescence staining (IIF). Scale bar, 100μm. I) Representative plots and summarized data of kidney immune cell phenotype in NCG-M HIS-Ctrl and NCG-M HIS-SLE group. Frequencies of CD69^+^ (activated) T and B cells were shown. J) Representative plots of CD69^+^(activated) B cells in human kidney biopsy sample. Data show mean ± SEM. Statistical analysis was performed using unpaired two-tailed *t*-test (A, G, H&I) and one-way ANOVA with Tukey multiple-comparison test (B, C, E&F). The exact *p* values are shown.

Taken together, these findings establish that TLR7 activation in NCG HIS mice: (1) drives potent human T cell responses, and (2) models SLE-associated immunophenotypes and renal pathology, while highlighting current limitations in B cell function that warrant further model optimization.

### TLR7 activation potentiates pathogenic effector B cell responses in next-generation NCG-M HIS mice

Recent advances in humanized mouse models have significantly improved human B cell maturation and class-switched antibody production. After screening several available strains — including NCG-X (NOD-*Prkdc^em26Cd52^Il2rg^em26Cd22^kit^em1Cin(V831M)^*/Gpt), NCG-X-TSLP (NOD-*Prkdc^em26Cd52^Il2rg^em26Cd22^kit^em1Cin(V831M)^Tg(mTSLP)918*/Gpt), NCG-Flt3-KO (NOD-*Prkdc^em26Cd52^Il2rg^em26Cd22^Flt3^em1Cd^*/Gpt), NCG-hIL15 (NOD-*Prkdc^em26Cd52^Il2rg^em26Cd22^Il15^em1Cin(hIL15)^*/Gpt) and NCG-M — we selected NCG-M mice for further studies. NCG-M mice are genetically modified to express low levels of human stem cell factor (SCF), granulocyte-macrophage colony-stimulating factor (GM-CSF), and interleukin-3 (IL-3) while simultaneously having their murine cytokine homologs knocked out. Compared to other strains tested, NCG-M HIS mice exhibited the highest baseline IgG levels and better peripheral B cell maturation (Figure 2B&2C). Further flow cytometry analysis revealed superior reconstitution of human T cells and dendritic cells in the spleen, as well as significant increases in all human immune cell numbers compared to NCG HIS mice (Figure S3a).

We then applied the TLR7 agonist administration protocol to NCG-M HIS mice and evaluated immune phenotype, autoantibody production, and kidney pathology (Figure 2D). Of note, the NCG-M HIS-SLE mice showed elevated anti-dsDNA antibody and augmented effector B cells in multiple tissues compared to both NCG-M HIS-Ctrl and NCG HIS-SLE mice (Figure 2E&2F, Figure S3b), showing a similar trend of the peripheral blood of lupus patients (Figure 2G). Fluorescent assays confirmed abundant anti-nuclear antibodies in the NCG-M HIS-SLE group (Figure 2H). Immunofluorescence staining of kidney sections revealed deposition of human IgG and mouse complement C3 in glomeruli, mirroring typical lupus nephritis (Figure S3c&3d). Interestingly, immune profiling in the kidney demonstrated heightened activation of both T and B cells (Figure 2I), similar to the immune landscape seen in lupus nephritis patients (Figure 2J).

Collectively, these results demonstrate that TLR7 agonist-induced NCG-M HIS mice robustly recapitulate key pathological features of human SLE, particularly the dysregulated B cell responses characteristic of the disease. Based on this superior phenotypic fidelity, we established NCG-M HIS mice as our primary experimental platform for subsequent investigations.

### Superior recapitulation of human SLE phenotypes by TLR7 agonist versus pristane in NCG-M HIS mice

Leveraging the superior human B cell-supportive environment of NCG-M HIS mice, we conducted a systematic comparison of lupus phenotypes induced by either the TLR7 agonist R848 or pristane — a historically employed lupus-inducing agent in humanized models. We evaluated serum autoantibodies, organ pathology, and immune dysregulation after two months of induction (Figure 3A). Compared to the pristane-induced-lupus (HIS-PIL) group, the R848-induced-lupus (HIS-RIL) group showed a significantly greater increase in total serum IgG levels (Figure S3a), accompanied by higher anti-nuclear antibody fluorescence intensity (Figure 3B&3C) and nearly twice the concentration of anti-dsDNA antibodies (Figure 3D). Consistent with TLR7 pathway activation, R848 induction markedly increased serum IFN-α, along with other disease-associated inflammatory cytokines (IL-6, IL-8, and IL-10; Figure 3E). Markers of kidney dysfunction, including elevated urea nitrogen and creatinine, also indicated more severe renal pathology in the HIS-RIL group (Figure 3F). Consistently, immunofluorescence staining revealed extensive IgM and IgG immune complex deposition throughout the glomeruli (Figure 3G) in the HIS-RIL group, compared to the HIS-PIL group. Beyond the kidneys, the HIS-RIL group developed similar degrees of interstitial lung fibrosis but displayed slightly greater liver inflammation than the HIS-PIL group (Figure S4b), indicating systemic involvement akin to human SLE.

**Figure 3.**
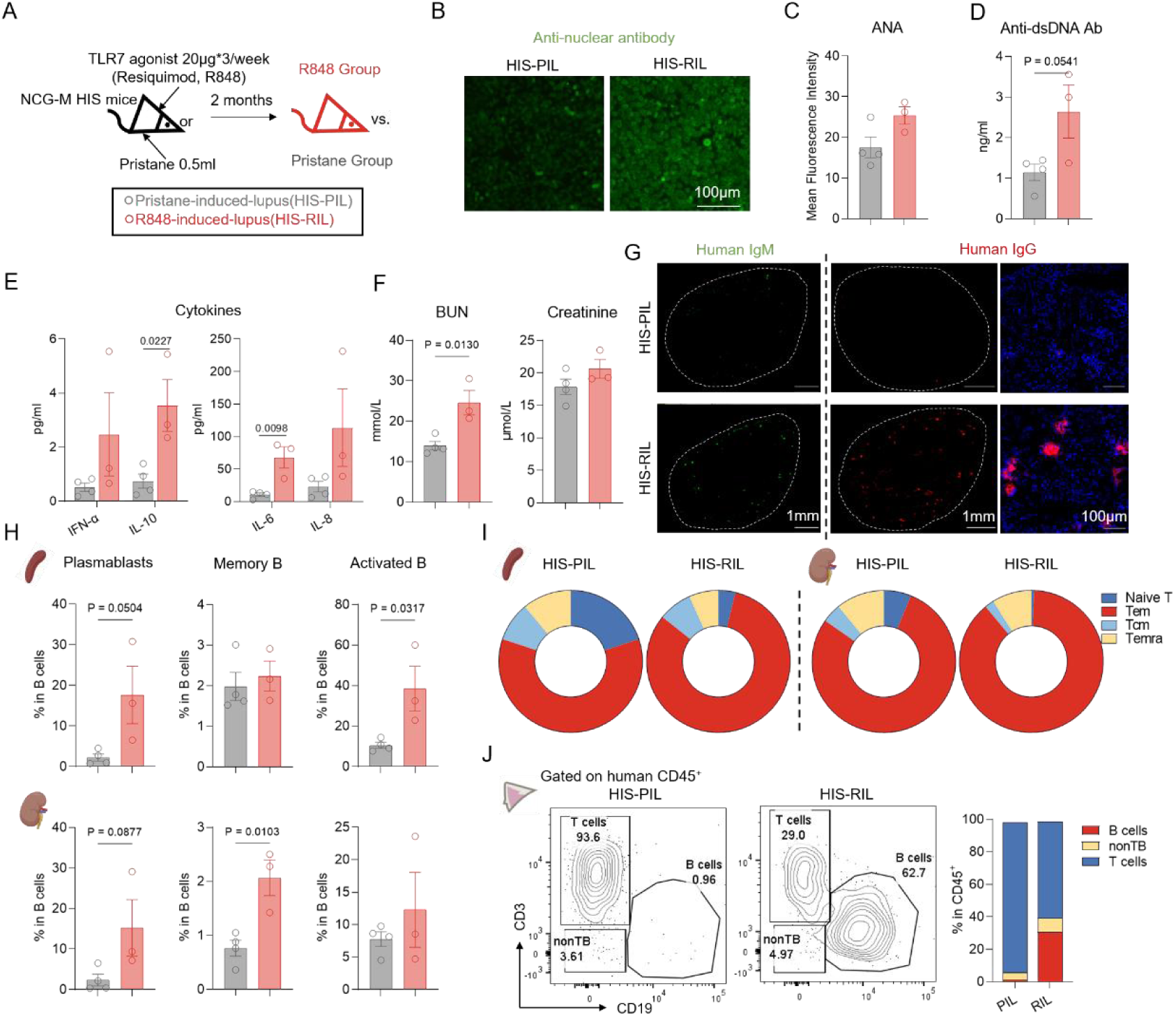
TLR7 agonist outperforms pristane in inducing lupus associated immune imbalance and tissue pathology. A) NCG-M HIS mice were either topically treated with TLR7 agonist (R848-induced-lupus group, HIS-RIL, *n=3*) or intraperitoneal injection with 500μl pristane (Pristane-induced-lupus group, HIS-PIL, *n=4*) for 8 weeks, disease phenotypes were determined at the endpoint. B-C) Representative images(B) and summarized data(C) of anti-nuclear antibody in serum from two groups detected by IIF. Scale bar, 100μm. D) Serum anti-dsDNA antibody levels detected by ELISA. E) Disease associated cytokine levels in serum detected by Legendplex. F) Urea nitrogen and creatinine level in serum from two groups detected by biochemical analysis. G) Immunofluorescence staining of human IgM (left, whole section) and human IgG (mid, whole section and right, enlarged version) deposition in kidneys from HIS-PIL and HIS-RIL group. Scare bar, 1mm or 100μm. H) Phenotypic analysis of B cells in spleen and kidney tissues from HIS-PIL and HIS-RIL group. Frequencies of CD24^−^CD38^hi^(plasmablast), CD38^−^CD27^+^ (memory) and CD86^+^(activated) cells were summarized. I) Phenotypic analysis of T cells in spleen and kidney tissues from HIS-PIL and HIS-RIL group. Frequencies of CD45RA^+^CCR7^+^ (naive), CD45RA^+^CCR7^−^ (Temra), CD45RA^−^CCR7^+^ (Tcm) and CD45RA^−^CCR7^−^ (Tem) cells were summarized. J) Phenotypic analysis of human immune cells in skin tissues from HIS-PIL and HIS-RIL group. Frequencies of CD3^+^ (T cells), CD19^+^ (B cells) and CD3^−^CD19^−^(nonTB) cells were summarized. Data show mean ± SEM. Statistical analysis was performed using unpaired two-tailed *t*-test. The exact *p* values are shown.

Flow cytometry analysis further demonstrated that B cells in the HIS-RIL group were more differentiated and activated, evidenced by higher frequencies of plasmablasts, memory B cells, and activated B cells infiltrating the spleen and kidney (Figure 3H, Figure S4c). Similarly, the HIS-RIL group exhibited increased, though not statistically significant, percentage of antibody-secreting cells in spleen and bone marrow (Figure S4d&4e). Human T cells also activated and differentiated towards the effector phenotype with a dramatic decrease in the naïve subset in the HIS-RIL model (Figure 3I, Figure S4f&4g). Notably, B cells were detected in the skin only in the HIS-RIL group (Figure 3J), a feature reported exclusively in the skin of SLE patients but absent in healthy donors.

In summary, our TLR7 agonist-induced NCG-M HIS-SLE mice closely recapitulates key immunological and pathological features of human SLE, and thus represents a transformative platform for preclinical evaluation of advanced SLE therapies.

### The HIS-SLE model enables comprehensive comparison of two B cell-targeted monoclonal antibodies

Monoclonal antibodies targeting B cells, notably rituximab and belimumab, are important therapeutic options for SLE patients refractory to conventional immunosuppressants. To establish proof-of-principle for preclinical utility of our HIS-SLE model, we assessed the efficacy of these two agents in HIS-SLE mice. Following antibody administration (Figure 4A), both treatments demonstrated reduced serum autoantibody levels compared to the untreated group (Figure 4B). Flow cytometry analysis revealed distinct immunomodulatory profiles reflecting each drug’s specific target. Rituximab efficiently depleted peripheral blood and splenic B cells, with residual populations enriched in CD20 negative plasma cells (Figure 4C, Figure S5a&5b). In bone marrow, rituximab depleted CD20^+^ B cells while sparing early B cell precursors (pro-B and pre-B cells), permitting rapid reconstitution that may limit long-term efficacy (Figure S5a). In comparison, belimumab exerted limited effects on total B cell counts while modestly increasing the proportion of effector B cell subsets (Figure 4C, Figure S5b), aligning with clinical observations that memory B cells and plasmablasts are relatively resistant to BAFF inhibition.^26,27^ Immunohistochemistry staining confirmed systemic CD20⁺ B cell clearance in non-lymphoid tissues (lung and liver) following rituximab treatment (Figure S5e).

**Figure 4.**
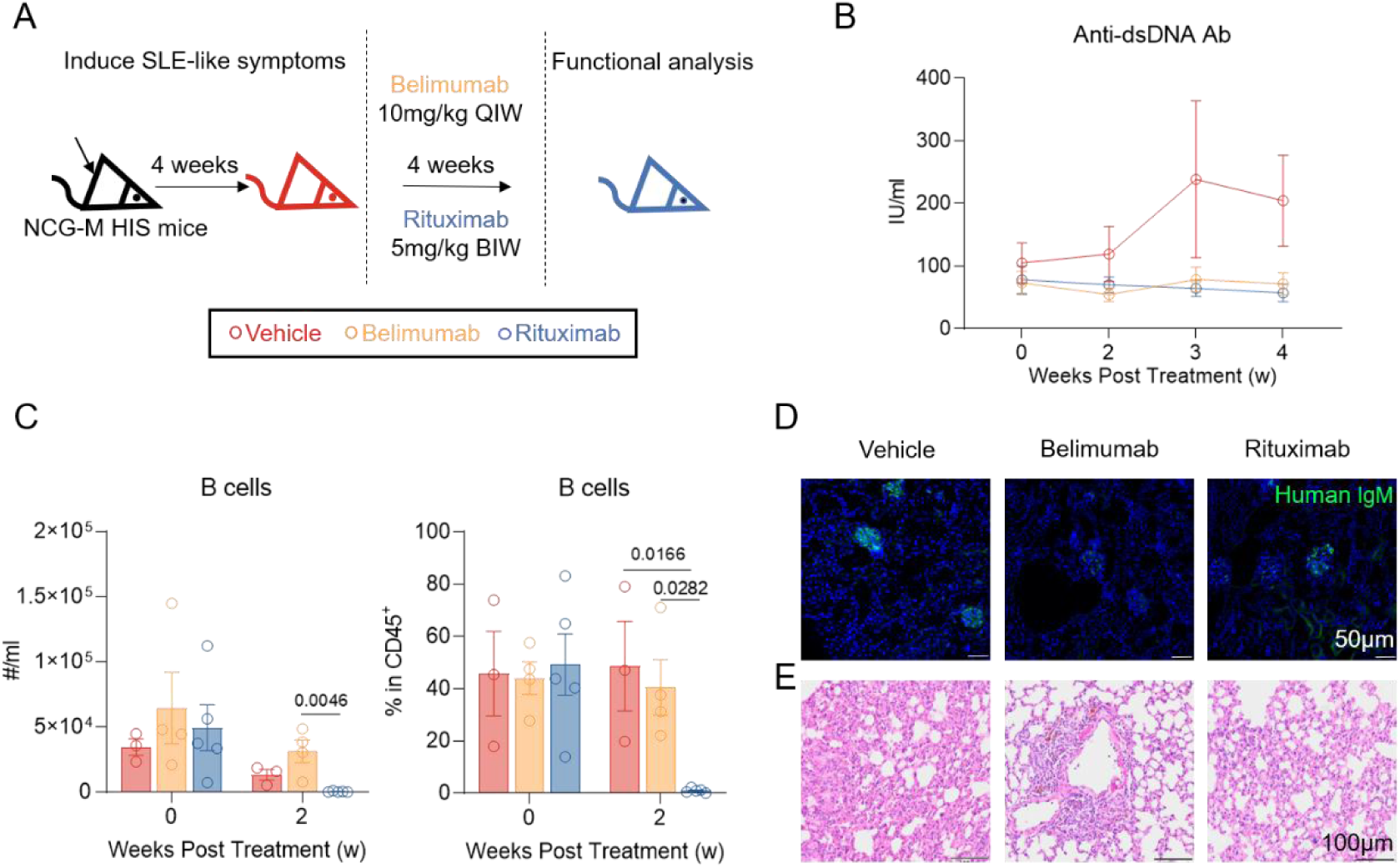
A comprehensive comparison of two B cell targeting monoclonal antibodies in HIS-SLE model. A) The schematic diagram of Belimumab and Rituximab *in vivo* treatment. TLR7 agonist-induced NCG-M HIS-SLE mice were either left untreated (Vehicle group) or intravenously injected with Belimumab or Rituximab at indicated doses and frequencies for four weeks. B) Anti-dsDNA antibody levels in serum from vehicle(*n=2*), Belimumab*(n=4)* and Rituximab*(n=5)* group at indicated time points detected by ELISA. C) Flowcytometry analysis of absolute numbers(left) and frequencies(right) of CD19^+^ B cells in peripheral blood at indicated time points. D) Immunofluorescence staining of human IgM deposition in the glomerular area of kidneys from Vehicle, Belimumab and Rituximab group. Scale bar, 50μm. E) HE staining of lung tissues from Vehicle, Belimumab and Rituximab group. Scale bar, 100μm. Data show mean ± SEM. Statistical analysis was performed using one-way ANOVA with Tukey multiple-comparison test. The exact *p* values are shown.

Given the interplay between B and T cells, we also investigated T cell differentiation. Notably, only belimumab promoted a shift from effector T cells to naïve T cells, suggesting improvement of the overall immune microenvironment (Figure S5c). Belimumab is also approved for lupus nephritis due to its renoprotective effects.^28^ Consistently, both antibodies reduced serum creatinine, urea nitrogen, and kidney IgG deposits (Figure S5d&5f), while belimumab showed slightly superior clearance of IgM deposits (Figure 4D). Histological evaluation of the lungs revealed fibrosis regression with both treatments (Figure 4E).

Our findings demonstrate that both rituximab and belimumab exhibit clinically meaningful efficacy in the HIS-SLE model, with rituximab achieving rapid peripheral B-cell depletion and belimumab modulating the immune microenvironment, while both treatments ameliorated serological abnormalities and renal pathology. However, the persistence of effector B-cell populations in protected tissue niches resulted in incomplete depletion and limited seroconversion, mirroring the sanctuary effects that contribute to clinical relapse patterns.^27^ These results highlight the need for next-generation approaches that can effectively penetrate tissue reservoirs and achieve durable, treatment-free remission.

### Preliminary evaluation of two T cell-engaged B cell-redirected regimens reveals distinct B cell elimination and reconstitution dynamics

Given the clinical promise of autologous CD19 CAR-T therapy in SLE and the emerging potential of CD3/CD19-targeted TCEs—both T cell-mediated, B cell-depletion strategies —we conducted a head-to-head evaluation of their safety and efficacy in HIS-SLE mice. Healthy donor T cells were positively selected, activated, and transduced with a lentiviral vector that contains the sequence for a single-chain variable fragment derived from an anti-human CD19 hybridoma clone (FMC63). Furthermore, the vector contains the information for a CD8-derived hinge and transmembrane domain, a CD3 ζ intracellular domain, and a 4-1BB costimulatory domain. Engineered T cells were then simultaneously knocked out for human leukocyte antigen (HLA)-A, HLA-B, HLA-C, and T cell receptor alpha constant (TRAC) by electroporation-based CRISPR Cas9 gene editing using Cas9 protein in complex with single guide RNAs (sgRNAs) for each target (Figure S6a). The final product showed a knock out efficiency of almost 100% and a CAR transduction efficacy of over 80% (Figure 5A). *Ex vivo*, UCAR-T cells showed efficient, dose-dependent B cell cytotoxicity: complete B cell elimination was achieved when effector-to-target ratios exceeded 1:2. In contrast, the bispecific antibody blinatumomab (targeting CD3 and CD19) failed to achieve full depletion, with nearly 20% of B cells persisting even at maximal concentrations (Figure S6b&6c).

**Figure 5.**
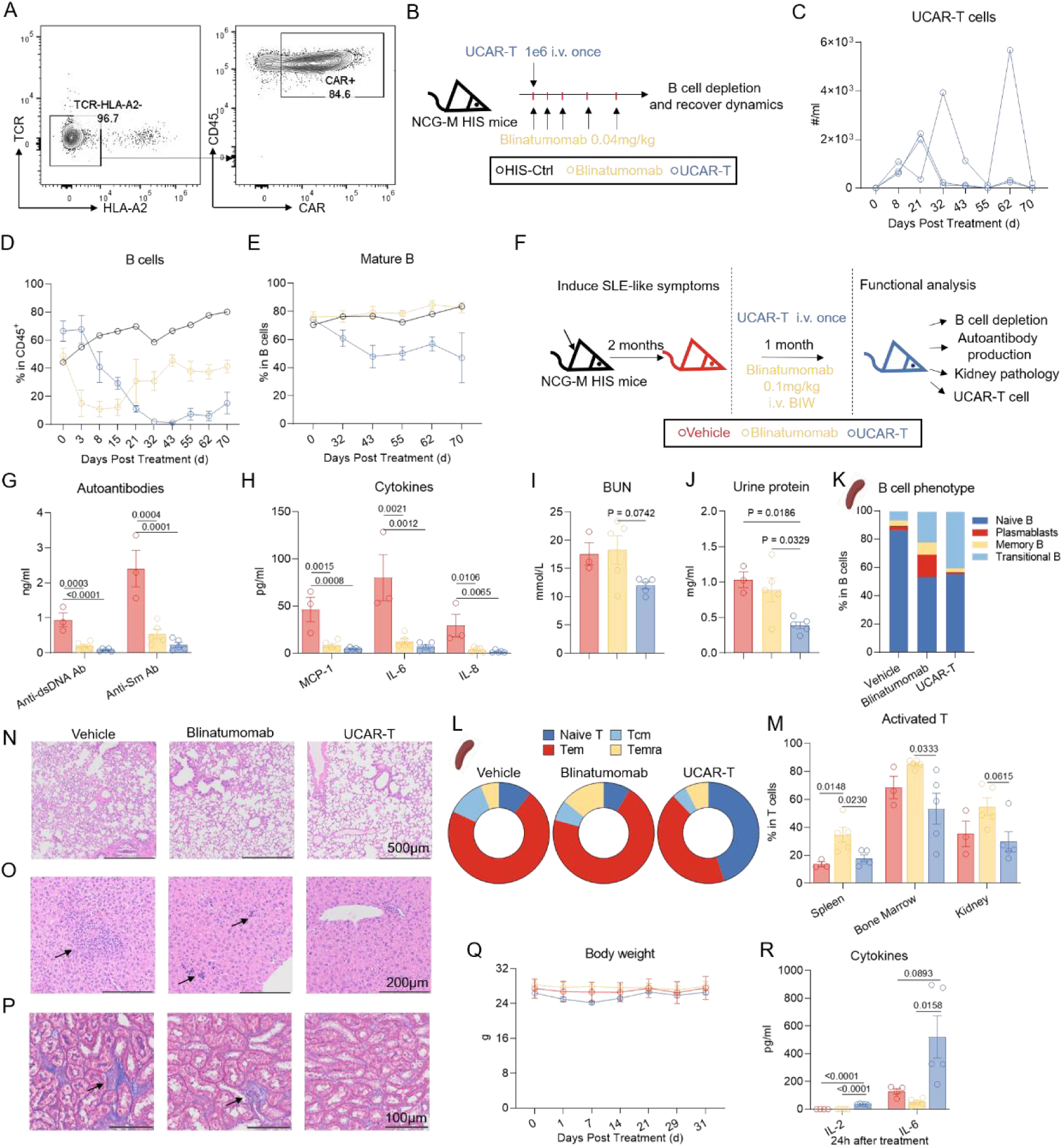
A comprehensive comparison of two T cell-mediated B cell targeting therapies in HIS-SLE model. A) Quality control of allogeneic CAR-T(UCAR-T) cells. Representative plots of TCR, HLA-A2 and CAR expression on UCAR-T cells were shown. B) The schematic diagram of UCAR-T cells and Blinatumomab in vivo application. Uninduced NCG-M HIS mice (HIS-Ctrl) were treated with either one million UCAR-T cells once(*n=3*) or T cell engager Blinatumomab at D0,1,2,4 and 6(*n=3*). The number of B cells in peripheral blood was monitored. C) Flowcytometry analysis of numbers of UCAR-T cells in peripheral blood at indicated time points. D) Flowcytometry analysis of frequencies of B cells in peripheral blood at indicated time points. E) Flowcytometry analysis of frequencies of mature B cells (CD24^int^CD38^int^) in peripheral blood at indicated time points. F) The schematic diagram of UCAR-T cells and Blinatumomab treatment in NCG-M HIS-SLE mice. NCG-M HIS-SLE mice induced with TLR7 agonist were treated with vehicle (Vehicle group, *n=3*), allogeneic CAR-T cells (UCAR-T group, *n=5*) or CD3-CD19 bispecific antibody (Blinatumomab group, *n=5*) for four weeks. Disease phenotypes were determined at the end point. G) Anti-dsDNA and anti-smith antibody levels in serum from three groups detected by ELISA. H) Disease associated cytokine levels in serum from three groups detected by Legendplex. I-J) Kidney function of three groups. Urea nitrogen levels in serum(I)and protein levels in urine(J) were summarized. K) Phenotypic analysis of B cells in spleen tissues from Vehicle, Blinatumomab and UCAR-T group. Frequencies of CD24^hi^CD38^hi^ (transitional B), IgD^+^CD27^−^ (naïve B), CD38^−^CD27^+^ (memory B) and CD24^−^CD38^hi^ (plasmablast) cells were summarized. L) Phenotypic analysis of T cells in spleen tissues from Vehicle, Blinatumomab and UCAR-T group. Frequencies of CD45RA^+^CCR7^+^ (naive), CD45RA^+^CCR7^−^ (Temra), CD45RA^−^CCR7^+^ (Tcm) and CD45RA^−^CCR7^−^ (Tem) cells were summarized. M) Flowcytometry analysis of frequencies of CD69^+^(activated) T cells in spleen, bone marrow and kidney tissues from three groups. N-P) Tissue pathology detected by HE staining and Masson Trichrome staining. (N) HE staining of lung tissues from three groups. Scare bar, 500μm. (O) HE staining of liver tissues from three groups. Arrows indicate the immune cells infiltrated area. Scare bar, 200μm. (P) Masson Trichrome staining of interstitial area of kidneys from three groups. Arrows indicate the fibrotic areas. Scare bar, 100μm. Q-R) Safety evaluation of the two therapies. (Q) Monitored body weight over the whole treatment period. (R) Cytokines in serum from three groups 24 hours after first injection. Data show mean ± SEM. Statistical analysis was performed using one-way ANOVA with Tukey multiple-comparison test. The exact *p* values are shown.

Next, we further explored the dynamics of B cell clearance and rebound after *in vivo* administration of allogeneic CAR-T cells or bispecific blinatumomab in NCG-M HIS mice (Figure 5B). One million UCAR-T cells were administered as a single intravenous infusion, whereas blinatumomab was given at 0.04 mg/kg—based on human-equivalent clinical dosing—and delivered in five separate infusions over seven days, for a cumulative dose of approximately 4μg. UCAR-T cells expanded rapidly post-infusion, peaking at days 21-32, during which time peripheral B cells were undetectable (Figure 5C&5D, Figure S6d). A second CAR-T cell expansion occurred around days 55-62, indicating the persistence of CAR-T cells and reactivation by reemergence of B cells. In contrast to the sustained B cell depletion by UCAR-T cells, blinatumomab displayed a transient effect characterized by rapid B cell clearance followed by swift reconstitution (Figure 5D). Analysis of mature and transitional B cell frequencies further confirmed the stronger clearance and repopulation control achieved by UCAR-T cells compared to blinatumomab (Figure 5E, Figure S6e&6f).

To assess the tissue-level clearance of B cells by CAR-T cells, we infused 5 million UCAR-T cells into HIS mice and monitored their expansion and distribution immediately after complete B cell clearance (Figure S6g). UCAR-T cells exhibited widespread infiltration across the spleen, bone marrow, kidney, liver, and lung, leading to efficient clearance of resident B cells (Figure S6h&6i). Notably, the most pronounced UCAR-T activation occurred in bone marrow, likely due to the high density of CD19 antigen (Figure S6j).

Collectively, these initial *in vitro* and *in vivo* assessments demonstrated that both interventions efficiently depleted B cells, albeit with distinct dynamics, potencies, and depths of depletion. Guided by these preliminary findings, we therefore refined the blinatumomab dosing regimen for subsequent studies in our HIS-SLE model.

### Head-to-head comparison of two T cell-engaged B cell-redirected regimens demonstrates superior efficacy of UCAR-T through profound depletion of tissue-resident B cells

After the functional validation of UCAR-T cells and blinatumomab, we moved forward to evaluate the safety and efficacy of those two B cell-targeted therapies in our HIS-SLE model (Figure 5F). UCAR-T cells were administered intravenously as a single dose of either 1 × 10^6^ or 5 × 10^6^ cells. Robust UCAR-T cell expansion drove rapid B cell elimination, rendering them undetectable by day7 — coinciding precisely with peak UCAR-T cell levels (Figure S7a&7b). As the initial blinatumomab dose was insufficient for B cell depletion (Figure 5D, Figure S6f), we escalated the dose and administered it twice weekly for four weeks to improve efficacy and offset its short half-life. Under this regimen, blinatumomab preserved its rapid clearance kinetics while preventing significant B cell recovery.

We next compared the effects of the two treatments on SLE pathology following sustained B cell depletion. Both therapies substantially lowered autoantibody titers and normalized inflammatory cytokines; CAR-T treatment was marginally more effective (Figure 5G&5H). More importantly, UCAR-T conferred significantly better renoprotection, with greater reductions in urine protein and serum urea nitrogen (Figure 5I&5J). Collectively, these results demonstrate improvement in key SLE serum markers by both treatments, with UCAR-T cells showing superior efficacy against lupus nephritis indicators.

Analysis of tissue-resident B cells further supported enhanced efficacy of UCAR-T cells over blinatumomab. B cells were nearly eradicated in bone marrow, spleen and kidney, with UCAR-T outperforming blinatumomab, especially in bone marrow (Figure S7c&7e). Concurrently, UCAR-T cells penetrated and persisted in these tissues for one month, confirming long-term survival (Figure S7g). Interestingly, the residual B cell phenotype in spleen differed substantially between groups: UCAR-T-treated mice had higher proportions of immature transitional B cells and fewer effector B cells (memory B cells and plasmablasts) compared to blinatumomab-treated mice, indicating a deeper reset of the B cell compartment (Figure 5K). Besides, IHC staining verified the depletion of B cells in the lung and liver, exhibiting systemic efficacy of the two regimens (Figure S7f).

Notably, our model accurately recapitulated how distinct B cell depletion strategies differentially remodeled the immune landscape, particularly affecting T cell populations. Blinatumomab activated T cells and promoted their differentiation toward effector phenotypes, driving antigen-specific cytotoxicity (Figure 5L&5M, Figure S7d). In contrast, UCAR-T not only eliminated B cells but also restored the T cell homeostasis by normalizing the over-activated and differentiated T cell phenotype, providing a more comprehensive and durable therapeutic effect (Figure 5L&5M, Figure S7d).

This improved immune homeostasis of UCAR-T was accompanied by amelioration of tissue pathology. Both blinatumomab and UCAR-T treatments led to a near absence of immune complex deposition in glomeruli, confirming renoprotection in HIS-SLE mice (Figure S7h), and both therapies comparably alleviated lung fibrosis (Figure 5N). Notably, UCAR-T conferred superior protection in the liver and interstitial area of the kidney: H&E staining demonstrated significantly attenuated hepatic inflammation (Figure 5O), while Masson staining showed greater reduction in renal interstitial fibrosis (Figure 5P) compared to blinatumomab.

### Safety assessment indicates a mild increase in UCAR-T toxicity yet preserved tolerability versus blinatumomab

Additionally, our model also enabled safety assessment of these novel B cell-targeted strategies, which is equally critical for therapeutic development. Cytokine release syndrome (CRS) and immune effector cell-associated neurotoxicity syndrome (ICANS) are commonly reported adverse events following CAR-T or TCE administration.^29^ In our study, we monitored cytokine profiles and body weight after the first injection of each therapy. Following CAR-T infusion, we observed increased IL-6 levels and a transient decrease in body weight, indicating a moderate but tolerable CRS that resolved quickly without intervention (Figure 5Q&5R). In contrast, the blinatumomab treated group maintained stable cytokine levels and body weight, demonstrating minimal toxicity and good tolerance. Overall, both regimens exhibited favorable safety profiles in our humanized mouse model.

Taken together, both UCAR-T and blinatumomab treatment displayed deep B cell eradication, leveraging excellent efficacy and controllable toxicity. UCAR-T cells achieved further reset of B cell population, reverse of T cell over-activation and protection of renal function, leading to more significant and durable effects.

## Discussion

In this study, we developed an advanced humanized mouse model of SLE that more accurately recapitulates patient phenotypes than previously available models. Building on this platform, we conducted head-to-head safety and efficacy evaluations of two monoclonal antibodies, and two T cell-mediated B cell-targeted therapies, generating valuable preclinical insights to guide clinical treatment decisions.

A fundamental challenge in modeling SLE using humanized mice lies in reconciling the essential role of pathogenic B cells in disease pathogenesis ^30^ with the inherent defects in human B cell development, maturation, differentiation, and class-switching commonly observed in these models.^31^ Strategies to improve B cell maturation and functionality in HIS mice including the introduction of human cytokines,^32^ promotion of secondary lymphoid tissue development,^33^ and dual humanization of both hematopoietic and organ-specific compartments.^34^ Herein, we employed the NCG-M model, genetically engineered to express human SCF, GM-CSF, and IL-3 at near-physiological levels, thereby promoting the expansion of human myeloid and CD4⁺ T cells^35^ and creating a supportive niche for human B cell maturation and class-switch recombination. This model has recently been applied to study tumor-infiltrating plasma cells and IgG secretion in glioblastoma,^36^ demonstrating pronounced plasma cell differentiation and high human IgG production. Following human HSC engraftment, NCG-M mice exhibit human-relevant B cell phenotypes and serum IgG profiles, alongside stable immune reconstitution and a prolonged lifespan (>20 weeks post humanization). These attributes make NCG-M HIS mice a highly suitable platform for modeling human SLE and preclinical therapeutic evaluation.

Pristane was ineffective in eliciting IFN-I signaling upregulation in previous humanized models, in contrast to its strong activity in conventional mice,^37^ most likely owing to insufficient human myeloid engraftment. ^36^We therefore conducted a head-to-head comparison of pristane and TLR7 agonist R848 in NCG-M HIS mice, revealing that R848 — consistent with its intrinsic mechanism — elicited significantly higher serum IFN-α levels and stronger systemic immune activation than pristane. Additionally, topical TLR7 agonist administration further recapitulated the cutaneous immune profile of SLE, mirroring the facial rash presentation observed in ∼50% of patients.^38^ Critically, TLR7 agonist induction offers enhanced experimental controllability with a shortened induction period, accelerating experimental timelines and ensuring utility for preclinical therapeutic studies.

Our model recapitulated clinical treatment failure and relapse patterns observed with monoclonal antibody therapies. Specifically, incomplete depletion of tissue-resident B cells was evidenced by persistent splenic plasmablast accumulation following rituximab treatment, mirroring clinical limitations.^39^ Conversely, post-belimumab expansion of BLyS-independent memory B cells and plasmablasts demonstrated BAFF pathway-independent escape mechanisms.^26,40^ Although neither rituximab nor belimumab achieved full seroconversion, reductions in autoantibody levels and glomerular immune complex burden confirmed the pathogenic role of B cells in lupus and the therapeutic value of B cell-directed regimens. Accordingly, future optimization should prioritize enhanced depletion of tissue-resident B cells and autoantibody-secreting plasmablasts through: combinatorial anti-CD20/anti-BAFF therapy to counter post-rituximab BAFF surges;^41–43^ Fc region-afucosylated antibodies (e.g., obinutuzumab) to enhance FcγRIII-mediated antibody-dependent cell-mediated cytotoxicity (ADCC);^44,45^ or direct plasma cell targeting via proteasome inhibitors (bortezomib)^46,47^ or anti-CD38 agents (daratumumab).^48,49^

CAR-T cells and TCEs represent the most rapidly advancing therapeutic modalities, offering potent cytotoxicity and enhanced tissue infiltration. Blinatumomab — the first clinically approved CD3 × CD19 bispecific TCE — lacks an Fc domain, conferring improved tissue penetration and safety at the expense of a short half-life. Our study provides the first safety-efficacy assessment of blinatumomab in a lupus-like human immune microenvironment. An initial short-course, low-dose regimen (extrapolated from rheumatoid arthritis protocols^13^) led to partial B cell depletion followed by rapid rebound, consistent with clinical outcomes in RA. Capitalizing on HIS model’s rapid feedback capacity, we subsequently implemented an extended 4-week oncology-style regimen. This approach achieved deeper and more sustained peripheral B cell clearance; however, residual B cells persisted within tissues, especially the bone marrow, and the proportion of splenic plasmablasts increased, recapitulating the incomplete lymphoid eradication observed in RA patients.^13,50^ Despite this, levels of autoantibodies and inflammatory markers declined significantly. However, chronic pathologies, including renal fibrosis and impaired kidney function, showed minimal resolution. While overall therapeutic efficacy was suboptimal, the favorable safety profile, as evidenced by relatively stable body weight and pro-inflammatory cytokine levels, suggests that blinatumomab may be better suited to patients with milder SLE activity and limited renal involvement.

Advancing beyond T cell engagers, UCAR-T cells exhibit superior tissue penetration to eradicate tissue-resident B cells and reset the B cell compartment, fundamentally remodeling the immune milieu. These findings align with clinical reports of CAR-T-induced B cell reconstitution^9,51^ while providing unprecedented multi-tissue analyses unobtainable in patients. Complete B cell clearance drove profound reductions in autoantibodies and inflammatory mediators, achieving seroconversion. Notably, UCAR-T treatment protected renal function and ameliorated renal fibrosis. Transient cytokine release and weight loss required no pharmacologic intervention, with no overt impact on animal well-being. Collectively, these data position UCAR-T therapy as a preferred intervention for severe SLE with renal involvement when supported by appropriate safety monitoring.

A recent pilot study compared the effects of various B cell – depleting strategies on lymphatic tissues and highlighted the superior tissue B-cell depletion achieved by CAR-T cells compared to three protein-based therapies, including rituximab and blinatumomab.^13^ Our HIS-SLE model not only corroborates these clinical findings but also transcends human biopsy limitations by enabling whole-body, multi-organ immune profiling and providing direct histopathological evidence of CAR-T tissue trafficking — addressing critical knowledge gaps where clinical studies can only infer penetration due to restricted tissue sampling diversity. Furthermore, the model circumvents major clinical trial challenges, such as variability introduced by heterogeneous patient populations,^52^ limited cohort sizes that complicate efficacy interpretation, and the high costs and operational complexity of human studies. Beyond enabling rigorous head-to-head comparisons, this system also supports rapid therapeutic optimization — as evidenced by the timely adjustment of blinatumomab dosing and treatment duration within weeks — thereby accelerating translational pipelines while conserving resources.

Given these strengths, the HIS-SLE model holds broad utility across multiple preclinical applications. It enables precise optimization of dosing regimens and treatment schedules prior to clinical translation, thereby enhancing trial efficiency and the probability of success. It also supports comprehensive, multi-tissue immunological and histopathological analyses, offering a powerful platform to validate clinical hypotheses or uncover novel mechanistic insights. Moreover, the model facilitates rapid, scalable, and prospective head-to-head comparisons of single agents and combination therapies, providing robust, translational evidence to guide personalized treatment strategies in SLE.

## Conclusion

Collectively, we have established and characterized an advanced humanized SLE mouse model — built on the NCG-M strain and driven by a short-course TLR7 agonist administration— that faithfully mirrors the immunopathology and therapeutic responses seen in patients. The head-to-head comparisons presented here demonstrate UCAR-T’s superior potential for comprehensive immune environment remodeling and durable remission achievement. Moreover, the model’s versatility opens new avenues for therapeutic optimization and mechanistic exploration, ultimately bridging the gap between preclinical findings and clinical application in managing refractory and severe SLE.

## Experimental Section

### Patients and Specimens

Peripheral blood mononuclear cells (PBMCs) were isolated from peripheral blood samples of systemic lupus erythematosus (SLE) patients and healthy controls (Tables S1, Supporting Information). Fresh kidney tissues from lupus nephritis (LN) patients who underwent renal biopsy at the Affiliated Huaian No.1 People’s Hospital of Nanjing Medical University (Jiangsu, China) were used for flow cytometry analysis. All SLE patients fulfilled the 1997 revised classification criteria of the American College of Rheumatology (ACR),^53^ and all LN patients met the ACR criteria for lupus nephritis.^54^ The medical data were extracted from the subjects’ electronic medical records. All participants provided the informed consent forms and this study was approved by the Ethical Committee of the Affiliated Huaian No.1 People’s Hospital of Nanjing Medical University (Approval no. KY-2022-068-01).

The human hematopoietic stem cells (HSCs) used in this study were obtained from Nanjing Drum Tower Hospital (Jiangsu, China), the Affiliated Hospital of Nanjing University Medical School. Prior to sample collection, informed consent was obtained from all patients, and the study was approved by the hospital’s ethics committee (Ethics Approval No. protocol #2021-488-01).

### Mouse models

Immunodeficient NCG (NOD-*Prkdc^em26Cd52^Il2rg^em26Cd22^*/Gpt, Strain NO. T001475), NCG-X (NOD-*Prkdc^em26Cd52^Il2rg^em26Cd22^kit^em1Cin(V831M)^*/Gpt, Strain NO. T003802), NCG-X-TSLP (NOD-*Prkdc^em26Cd52^Il2rg^em26Cd22^kit^em1Cin(V831M)^Tg(mTSLP)918*/Gpt, Strain NO. T050142), NCG-Flt3-KO (NOD-*Prkdc^em26Cd52^Il2rg^em26Cd22^Flt3^em1Cd^*/Gpt), NCG-hIL15 (NOD-*Prkdc^em26Cd52^Il2rg^em26Cd22^Il15^em1Cin(hIL15)^*/Gpt, Strain NO. T004886) and NCG-M (NOD-*Prkdc^em26Cd52^Il2rg^em26Cd22^Rosa26^em1Cin(hCSF2&IL3&KITLG)^*/Gpt, Strain NO. T036669) strains were obtained from Gempharmatech (Nanjing, China) and reconstructed with human immune system as described below. All animals were housed under specific pathogen-free conditions, and all experiments were performed with the approval and oversight of the Institutional Animal Care and Use Committee (IACUC) at the Model Animal Research Center in Nanjing University (AP:LY-02).

### Human Immune System (HIS) mouse model

NCG or NCG-M HIS mice were reconstituted as described in this protocol^23^. Briefly, human fetal liver CD34^+^ cells were isolated using microbead kit (130-046-703, Miltenyi Biotec) and subsequently phenotyped for CD38 expression by flow cytometry. Newborn NCG-M pups (4-6 days old) received sublethal irradiation (80cGy) before intrahepatic injection of 5×10^4^ CD34^+^CD38^−^ human fetal liver HSCs. After 12 weeks, 50μL blood was drawn from each mouse to check human cell reconstitution. NCG or NCG-M HIS mice with more than 2 × 10^5^/ml human CD45^+^ cells in the blood were used for subsequent experiments.

### Humanized SLE mouse model

Experimental mice were chosen randomly, regardless of sex and reconstitution level. We performed the topical induction regimen following the method in this paper with a reduction in the dose.^21^ Briefly, the skin on the right ears of the mice were treated topically, 3 times weekly, with 20μg of resiquimod (R848; HY-13740, MCE) in 50μl of acetone. Mice treated with acetone only served as solvent control. Blood was drawn from facial vein every two weeks for flowcytometry analysis and plasma collection. 6-8 weeks later, mice were either sacrificed for end-point disease symptoms evaluation or given treatments for efficacy evaluation. For previously constructed pristane induced lupus model^18^, 12–13 weeks old NCG-M HIS mice were injected with 500μl pristane (1921-70-6, Sigma Aldrich) and waited for 6-8 weeks for the development and progression of disease.

### Generation of allogeneic anti-CD19 chimeric antigen receptor (CAR)-T cells

The CD19 CAR sequence was cloned into the lentiviral vector backbone. T cells collected from healthy donors were subjected to enrichment with magnetic separation using anti-CD3 microbeads (Miltenyi Biotec) and activation with T Cell TransAct (Miltenyi Biotec). 24 h after activation, lentivirus was mixed with T cells at a final dilution of MOI 3 in 6 well round bottom plates. Per well of the 6 well plate, 5 ×10^6^ T cells were mixed with media containing lentivirus (1 × 10^9^ TU/ml) in 15 μ l and then spinfected at 800 g for 60 min. Cells of the same condition were pooled and seeded at 5×10^5^ cells/ml in cell culture flask. 2 days post transduction, cells were collected and pelleted in preparation for nucleofection of Cas9 ribonucleoproteins (RNPs) to generate TCR, HLA-I knockouts. RNPs for each target locus were complexed separately at a 1:2 sgRNA (GenScript Biotech): SpCas9 (Thermo) ratio, followed by mixture of the two RNPs at 42pmol TRAC RNP, 42pmol HLA-I RNP per million cells, The RNP mixture was diluted in P3 solution (Lonza). mixed with the cell pellet, and nucleofected using the EO115 program (Lonza). Nucleofected cells were seeded at 1 ×10^6^ cells/ml, a magnetic separation using anti-CD3 microbeads (Miltenyi Biotec) was conducted to remove CD3^+^ cells to improve purity of allogeneic universal CAR-T cells. T cells were cultured in X-VIVO 15 medium (Lonza) supplemented with 5% CTS Immune Cell Serum Replacement (Gibco), recombinant human IL-2 (200 U/ml), IL-7 (10 ng/ml) and IL-15 (5 ng/ml) for 14 days. Cell products were stored in cryoprotectant after harvest.

### In vitro cytotoxicity assay

PBMCs from healthy donors were enriched by Ficoll (17544203, Cytiva) density gradient centrifugation and the percentage of B cells were calculated by flowcytometry analysis. PBMCs were either co-cultured with allogeneic CAR-T cells at an E:T (effector: target=CAR-T cells: calculated B cells) ratio from 4:1 to 1:8 or cultured with different concentrations of Blinatumomab for 24 hours. After the incubation, cells were prepared for and analyzed by flow cytometry. B, T and CAR-T cells were differentiated by CD19/CD3/CAR staining. The percentage of living CD19^+^ B cells was determined and used to calculate the percent cytotoxicity by the following equation: % Cytolysis = (% B Cells (PBMC alone) - % B Cells (Sample of interest)) / % B Cells (PBMC alone) ×100.

### In vivo monoclonal antibody, CAR-T and T cell engager treatment protocols

All therapeutic agents were administered intravenously with the following regimens: Belimumab (10 mg/kg once weekly), Rituximab (5 mg/kg twice weekly on days 0 and 3), and Blinatumomab (0.1 mg/kg twice weekly on days 0 and 3). For CAR-T therapy, cryopreserved cells were thawed and infused as a single dose on day 0, with low- and high-dose groups receiving 1×10⁶ or 5×10^6^ cells, respectively. Mice were monitored carefully for body weight fluctuations and fur condition throughout the 28-day treatment period, with the experimental endpoint defined as 30 days post-treatment initiation.

### Flow cytometry

Spleen, lymph node and liver tissues were grinded with a syringe plunger and filtered through a 70μm cell strainer. Bone marrow cells were crushed from dissected femurs and tibias into complete medium, gently pipetted and filtered through a 100μm nylon. Kidney and lung tissues were minced and digested with 40μg/ml DNase I and 1mg/ml collagenase D for 30–45 min at 37 °C. Cells were filtered through a 40μm strainer, suspended in 40% Percoll underlaid with 80% Percoll (GE Healthcare Life Sciences, UK) and centrifuged. The middle layer, an enriched population of leukocytes, was harvested. The enriched immune cells were than washed and resuspended in staining buffer consisting of a live/dead dye (eFluor™ 450/eFluor™ 780) and IgG from human serum for 10-15min to exclude dead cells and prevent unspecific staining. After being washed with staining buffer once, cells were then incubated with a mixture of primary antibodies. Alternatively, following surface staining, cells were incubated with fixation-permeabilization buffer, washed with permeabilization buffer (Fixation/Permeabilization Solution Kit; BD Biosciences, CA, USA), and then incubated with antibodies against intracellular antigens. For the cytokine secretion assay, cells were stimulated with Phorbol 12-myristate 13-acetate (PMA) in combination with ionomycin for four hours in the incubator. The staining protocol was similar following the process of surface staining and intracellular staining using another kit (Fixation/Permeabilization Kit, Invitrogen™ eBioscience™, USA). Samples were processed by a NovoCyte Flow cytometer system (Agilent Technologies Inc., USA) and analysed with FlowJo software (FlowJo, LLC, OR, USA). The antibodies used for flow cytometry are listed in Supplementary Table 2.

### Serum cytokine level by Legendplex

Peripheral blood from mice and human were collected by centrifuge at 2000g for 10 minutes. We performed multiplexed detection of cytokine production using the LEGENDplex™ Human Essential Immune Response Panel (13-plex) (740930, Biolegend) according to manufacturer’s instructions.

### Serum antibody profile by Enzyme-linked immunosorbent assay (ELISA)

Levels of anti-dsDNA antibodies in HIS mice serum were quantified using ELISA kits designed for human anti-dsDNA antibody (CB13357-Hu, COIBO BIO). Levels of anti-smith antibodies in humanized mouse serum were quantified using ELISA kits designed for human anti-smith antibody (CB19966-Hu, COIBO BIO). Human IgG in the serum of HIS mice was performed by coating plates with Goat Anti-Human IgA+IgG+IgM (H+L) (109-005-064, Jackson ImmunoResearch) and detecting with HRP-conjugated Goat Anti-Human IgG, Fcγ fragment specific (109-005-008, Jackson ImmunoResearch). All the measurements were conducted following the manufacturer’s instructions. In addition, urine protein was detected by using protein detection kits (KGA804, KeyGen Biotech).

### Indirect immunofluorescence staining (IIF) staining of Anti-nuclear Antibody (ANA)

IFAs were performed as described previously.^55^ In brief, HEp-2 cell coated slides (4220-12CN, MBL Life Science) were incubated at room temperature with serum from HIS mice for 30min, washed in PBS and visualized with FITC anti-human IgG by fluorescence microscopy. Controls included positive and negative serum from SLE patients and healthy donors.

### Histological analysis

Kidney, liver, lung and brain tissues were collected at the endpoint, fixed in 4% paraformaldehyde (P0099, Beyotime Biotechnology), embedded in paraffin, sectioned into 5 µm-thick, and stained for hematoxylin and eosin (H&E) and Masson trichrome. Slides were read with an Olympus microscope (Olympus VS200 Slide Scanner).

For immunohistochemistry, tissues were fixed in 4% paraformaldehyde, embedded in paraffin, sectioned into 5µm-thick, and stained with rabbit anti-human CD20(A4893, ABclonal). An anti-rabbit HRP (111-035-003, Jackson Immuno Research) was used as secondary antibody. Images were acquired using Olympus microscope (Olympus VS200 Slide Scanner) and analyzed by VS200 ASW software (Olympus).

For immunofluorescence evaluation of IgM, IgG and C3 deposits, kidneys were flash-frozen in optimal cutting temperature (OCT) compound (23-730-571, Fisher Biotec) and sectioned at 5μm thickness on a cryotome (Leica Biosystems). Frozen sections were thawed, blocked in blocking buffer, and incubated with Cy5 conjugated goat anti-human IgG (ab97172, Abcam), FITC conjugated mouse anti-human IgM(314506, Biolegend) or rabbit anti-mouse complement3(ab11862, Abcam) at 4°C overnight. Alternatively, slides stained with anti-mouse C3 were further incubated with AF647 goat anti-rat IgG (112-625-003, Jackson Immuno Research) at room temperature for 1 hour. All slides were mounted with DAPI (HY-D0814, MCE) and visualized using an Olympus microscope (Olympus VS200 Slide Scanner). The level of glomerular fluorescence from fluorescence microscopy images was measured by ImageJ software.

### Biochemical Tests

HIS mice were euthanized and plasma was collected through centrifugation of peripheral blood. Level of various biochemical indicators were analyzed by automated BX3010 chemistry analyzer (Sysmex) according to the manufacturer’s instruction

### Statistical Analysis

Data for all experiments were analyzed with GraphPad Prism 10 statistical software (San Diego, USA). Statistical significance was determined using unpaired two-tailed Student’s t test. When more than two groups of samples were compared, one-way ANOVA (with Tukey’s multiple-comparison post-tests) was used. Graphs containing error bars show means ± SEM. If not specially stated, *p* values less than 0.05 were considered statistically significant (\**p*< 0.05, \*\**p*< 0.01, \*\*\**p*< 0.001, \*\*\*\**p*< 0.0001). The exact *p* values are shown for each data.

## Author Contributions

L.S., Y.L. and R.Z. designed the study. L.S., Y.L., S.D. and S.L. supervised the study and revised the manuscript. R.Z., S.D. and J.L. conducted the investigation process. D.R. and J.F. contributed to methodology design and analysis of data. S.L., Z.Z., C.C. and S.Y. performed the animal experiments and conducted data collection. K.W., Y.C. and Y.H. managed the patient resources. X.S. provided the anti-CD19 allogeneic CAR-T cells. All authors read and approved the manuscript.

## Acknowledgements

This study was supported by grants from the Key Program of the National Natural Science Foundation of China NO.82330055 (L.S.), National Key Research and Development Program of China NO.2020YFA0710800 (L.S.), Joint Fund of the National Natural Science Foundation of China (U24A20380) (L.S.), Key Projects of the Jiangsu Basic Research Program (BK20243061) (L.S.), Scientific and Technological Innovation 2030 NO. 2023ZD0500404 (Y.L.), National Natural Science Foundation of China NO.32122035 (Y.L.), 32471000 (Y.L.), 82001717 (S.D.) and Science and Technology Innovation Key R&D Program of Chongqing (CSTB2024TIAD-STX0001, X.S.).

## Data Availability Statement

All data generated or analyzed during this study are available from the corresponding authors upon reasonable request.

## Supporting Information

**Figure S1 (related to Figure 1):**
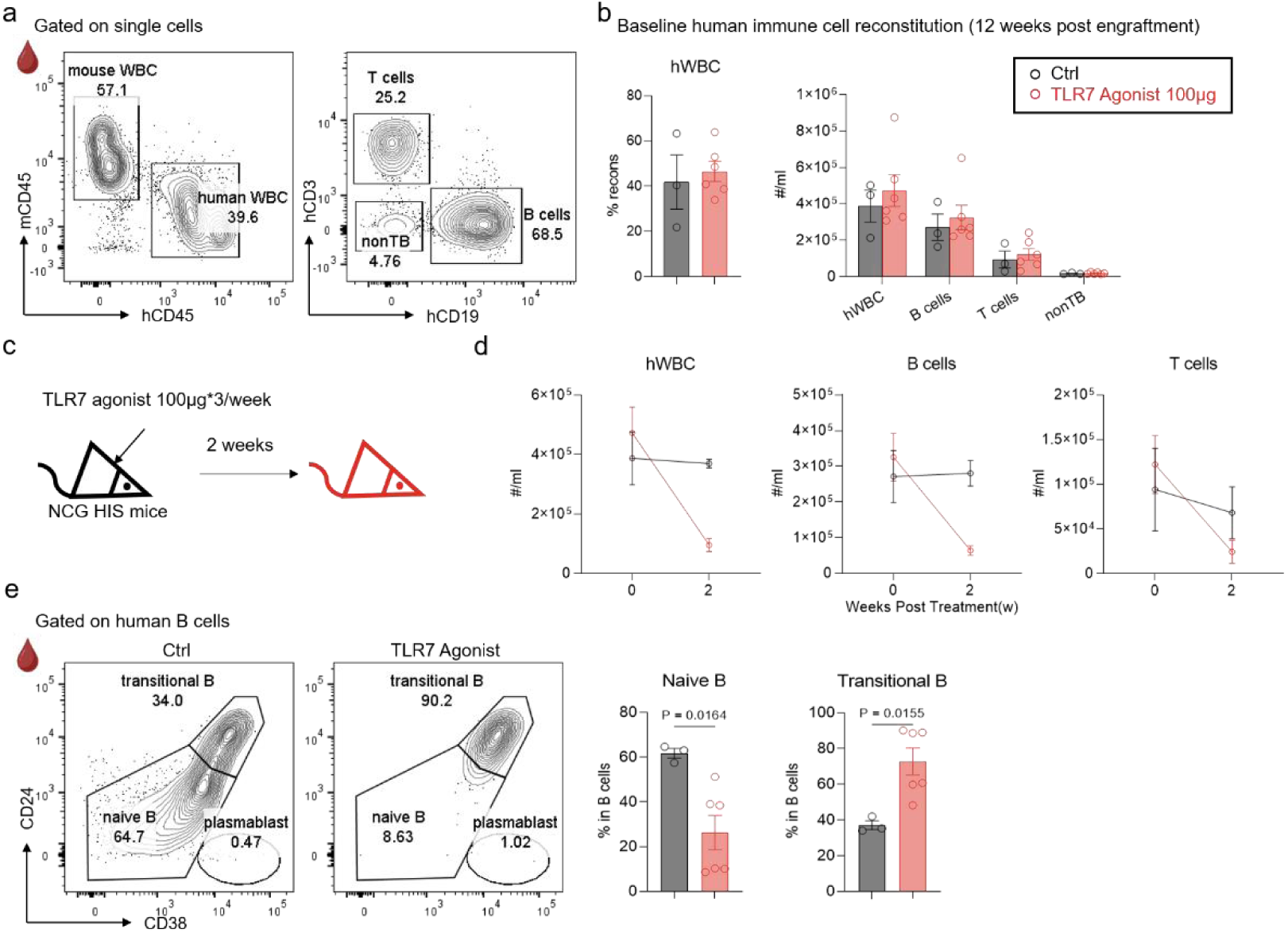
Common dose TLR7 agonist induction resulted in dramatic human cell loss. (a) Representative plots of human immune cell reconstitution 12 weeks post HSC transfer in NCG mice. (b) Summarized data of the percentage of human immune cells in total WBCs and the absolute numbers of different cell types. (c) NCG HIS mice were topically treated with common dose(100μg*3/week) TLR7 agonist for 2 weeks. Ctrl group, *n=3*. TLR7 agonist (100μg)-induced group, *n=6*. (d) Flowcytometry analysis of numbers of human cells at indicated time points. (e) Phenotypic analysis of B cells in peripheral blood from two groups. Frequencies of CD24^hi^CD38^hi^(transitional) and CD24^int^CD38^int^(naive) cells were summarized. Data show mean ± SEM. Statistical analysis was performed using unpaired two-tailed *t*-test. The exact *p* values are shown.

**Figure S2 (related to Figure 1):**
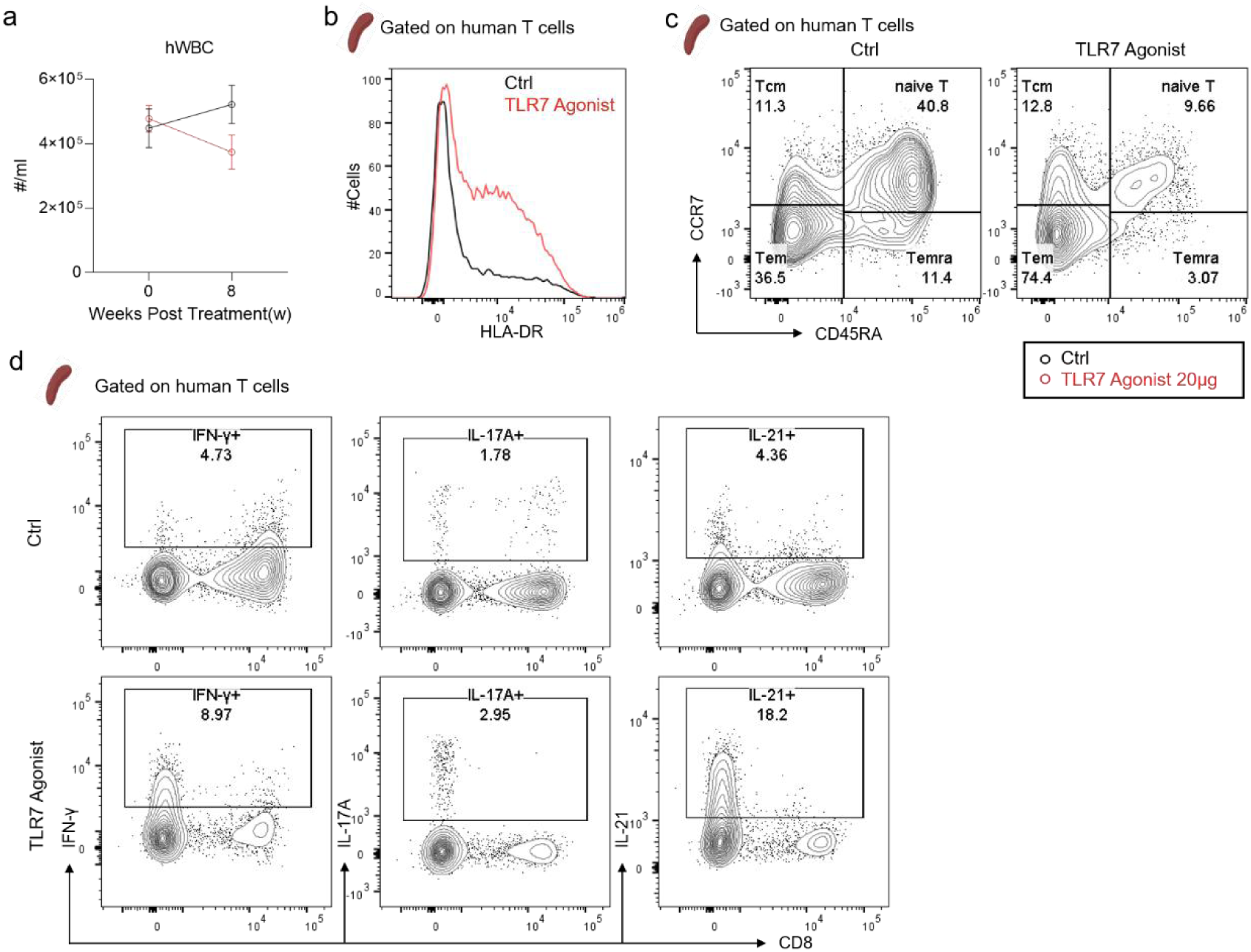
(a) Flowcytometry analysis of numbers of human cells at indicated time points. (b-c) Representative plots depicting HLA-DR (b), CD45RA and CCR7 (c) expression on human T cells from spleens of NCG HIS mice: Ctrl vs. TLR7 agonist-induced group (related to Figure 1C). (d) Representative plots of several cytokines expression on human T cells from spleens of the indicated two groups (related to Figure 1E). Data show mean ± SEM.

**Figure S3 (related to Figure 2):**
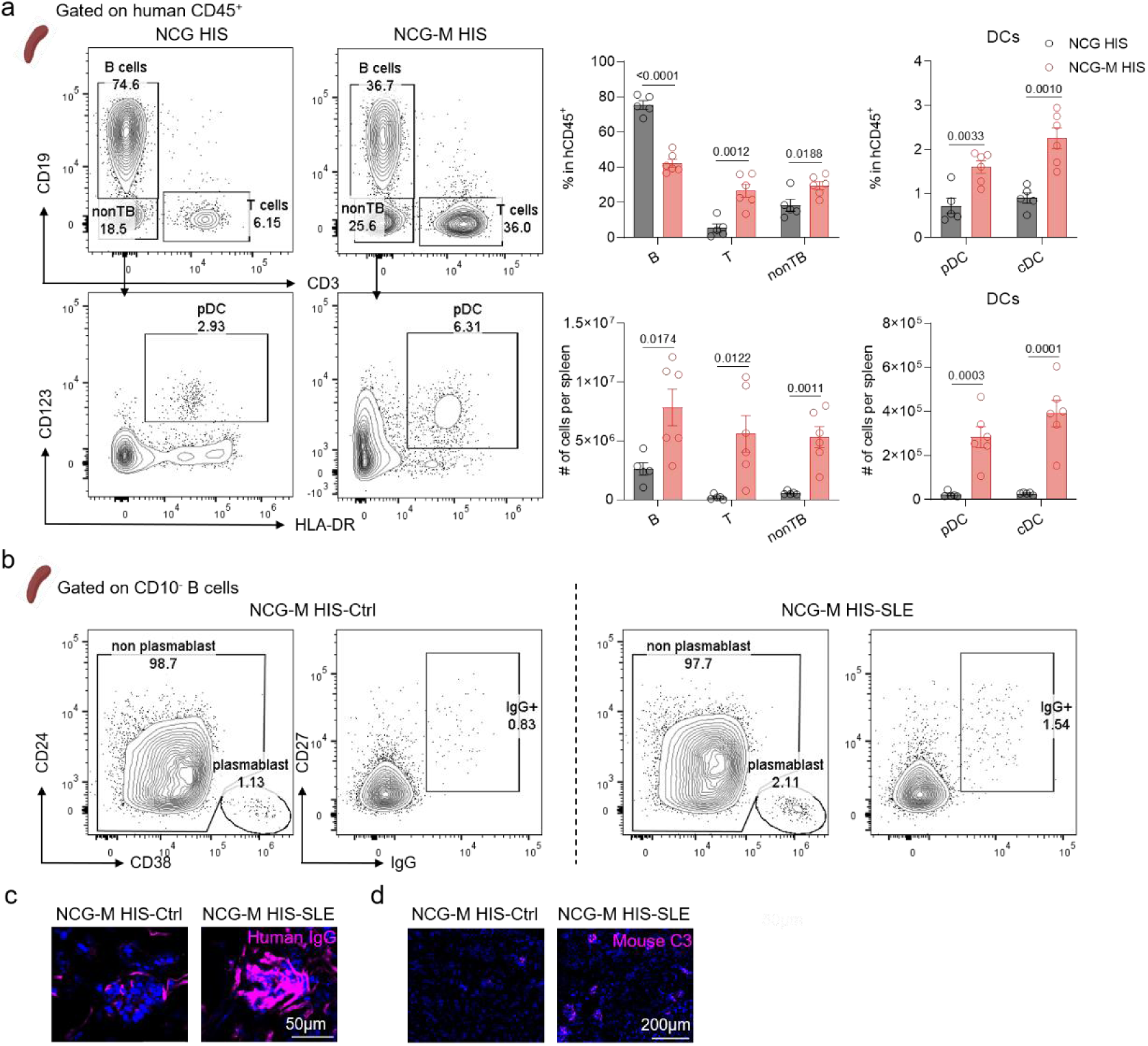
Screening identifies NCG-M HIS mice with enhanced human immune system reconstitution for SLE modeling. (a) Phenotypic analysis of human immune cells in spleens from NCG HIS*(n=5)* and NCG-M HIS mice*(n=6)*. Frequencies and absolute numbers of CD19^+^ (B cells), CD3^+^(T cells), CD19^−^CD3^−^(nonTB), CD19^−^CD3^−^HLA-DR^+^CD123^+^(pDC) and CD19^−^CD3^−^HLA-DR^+^CD1c^+^(cDC) cells were summarized. (b) Representative plots of Figure 2F. (c-d) Immunofluorescence staining of human IgG (c) and mouse complement3(C3, d) deposition in glomerular area of kidneys from NCG-M HIS-Ctrl and NCG-M HIS-SLE group. Scare bar, 50 or 200μm.Data show mean ± SEM. Statistical analysis was performed using unpaired two-tailed *t*-test. The exact *p* values are shown.

**Figure S4 (related to Figure 3):**
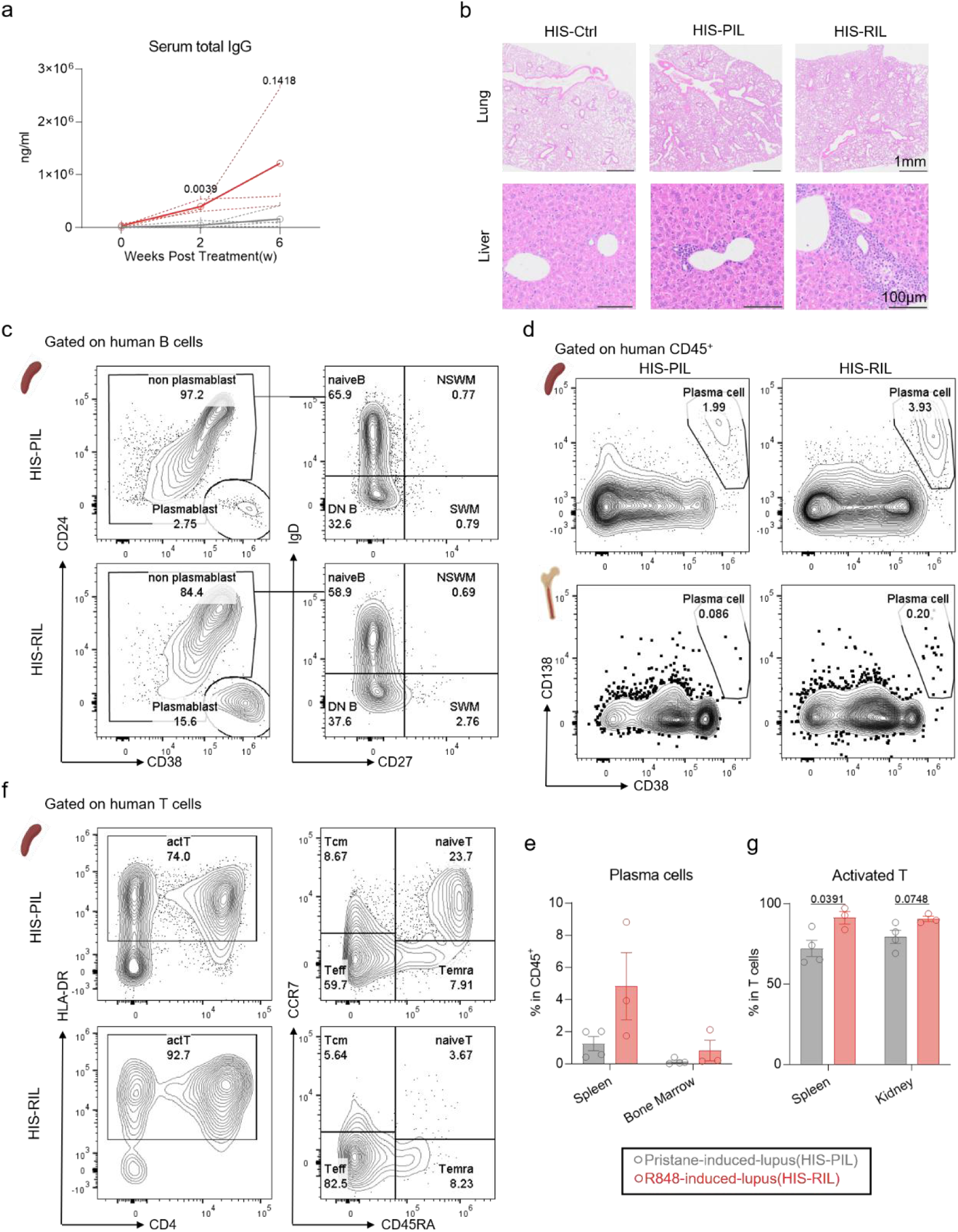
(a) Total IgG level in serum at indicated time points from HIS-PIL*(n=4)* and HIS-RIL*(n=3)* group detected by ELISA. Solid lines depicted the average level of each group and dotted lines showed the level of each individual. (b) Representative HE-staining images of lung(up) and liver(down) from HIS-Ctrl, HIS-PIL and HIS-RIL group. (c) Representative plots of B cell subsets in spleens from HIS-PIL and HIS-RIL group (related to Figure 3H). (d-e) Representative plots(d) and summarized data(e) of frequencies of CD38^hi^ CD138^+^ plasma cell in spleen and bone marrow tissues from HIS-PIL*(n=4)* and HIS-RIL*(n=3)* group. (f) Representative plots of T cell subsets in spleens from HIS-PIL and HIS-RIL group (related to Figure 3I). (g) Summarized data of frequencies of HLA-DR^+^ (activated) T cells in spleen and kidney tissues. Data show mean ± SEM. Statistical analysis was performed using unpaired two-tailed *t*-test. The exact *p* values are shown.

**Figure S5 (related to Figure 4):**
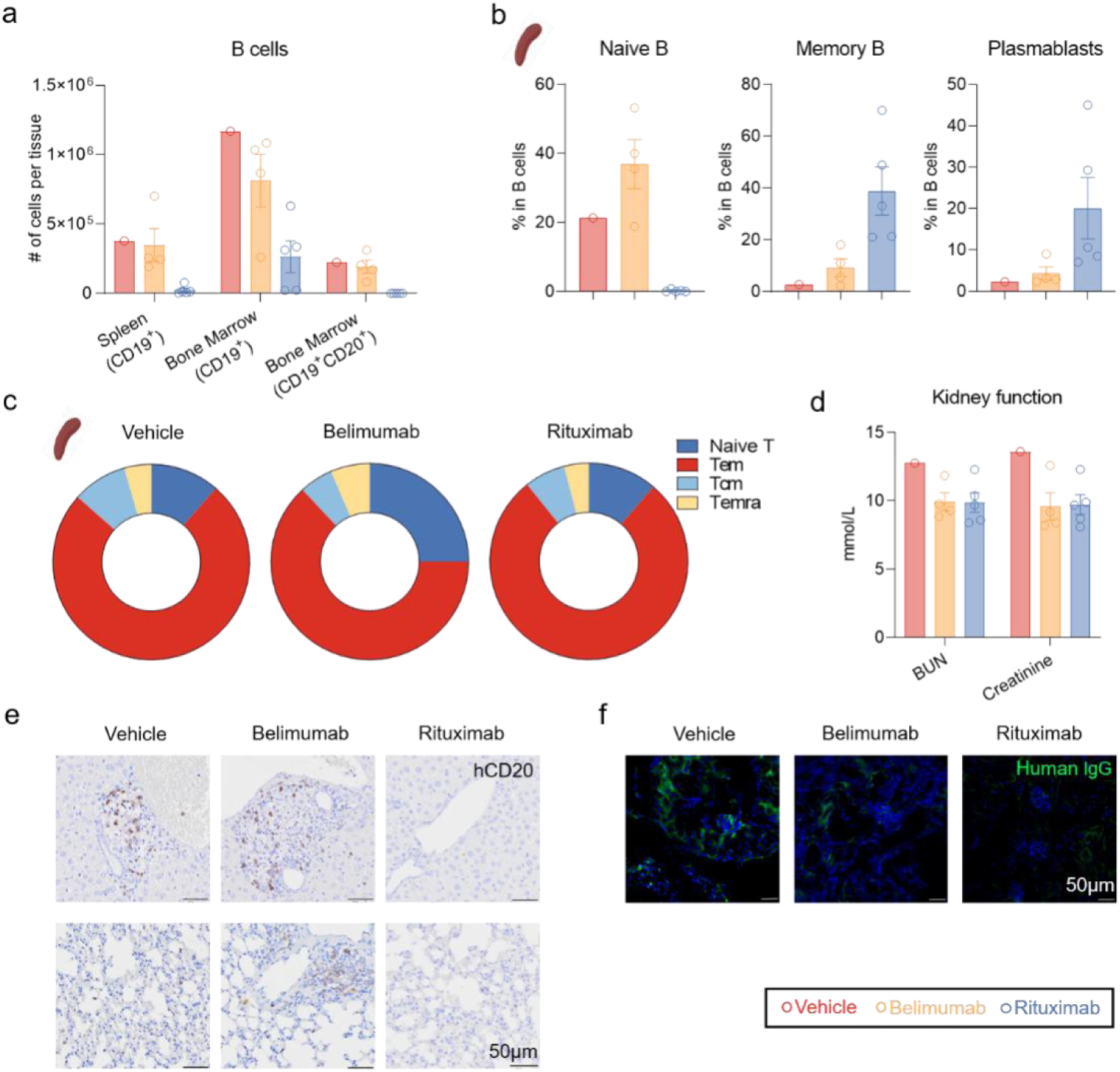
(a) Flowcytometry analysis of absolute numbers of CD19^+^ or CD19^+^CD20^+^B cells in tissues from Vehicle*(n=1)*, Belimumab*(n=4)* and Rituximab*(n=5)* group. (b) Phenotypic analysis of B cells in spleens from Vehicle, Belimumab and Rituximab group. Frequencies of CD24^int^CD38^int^ (naive), CD38^−^CD27^+^ (memory) and CD24^−^CD38^hi^ (plasmablast) cells were summarized. (c) Phenotypic analysis of T cells in spleens from three groups. Frequencies of CD45RA^+^CCR7^+^ (naive), CD45RA^+^CCR7^−^ (Temra), CD45RA^−^CCR7^+^ (Tcm) and CD45RA^−^CCR7^−^ (Tem) cells were summarized. (d) Urea nitrogen (BUN) and creatinine levels in serum from three groups detected by biochemical analysis. (e) Immunohistochemistry staining of human CD20 in the liver(up) and lung(down) tissues from Vehicle, Belimumab and Rituximab group. Scale bar, 50μm. (f) Immunofluorescence staining of human IgG deposition in the glomerular area of kidneys from three groups. Scale bar, 50μm. Data show mean ± SEM.

**Figure S6(related to Figure 5):**
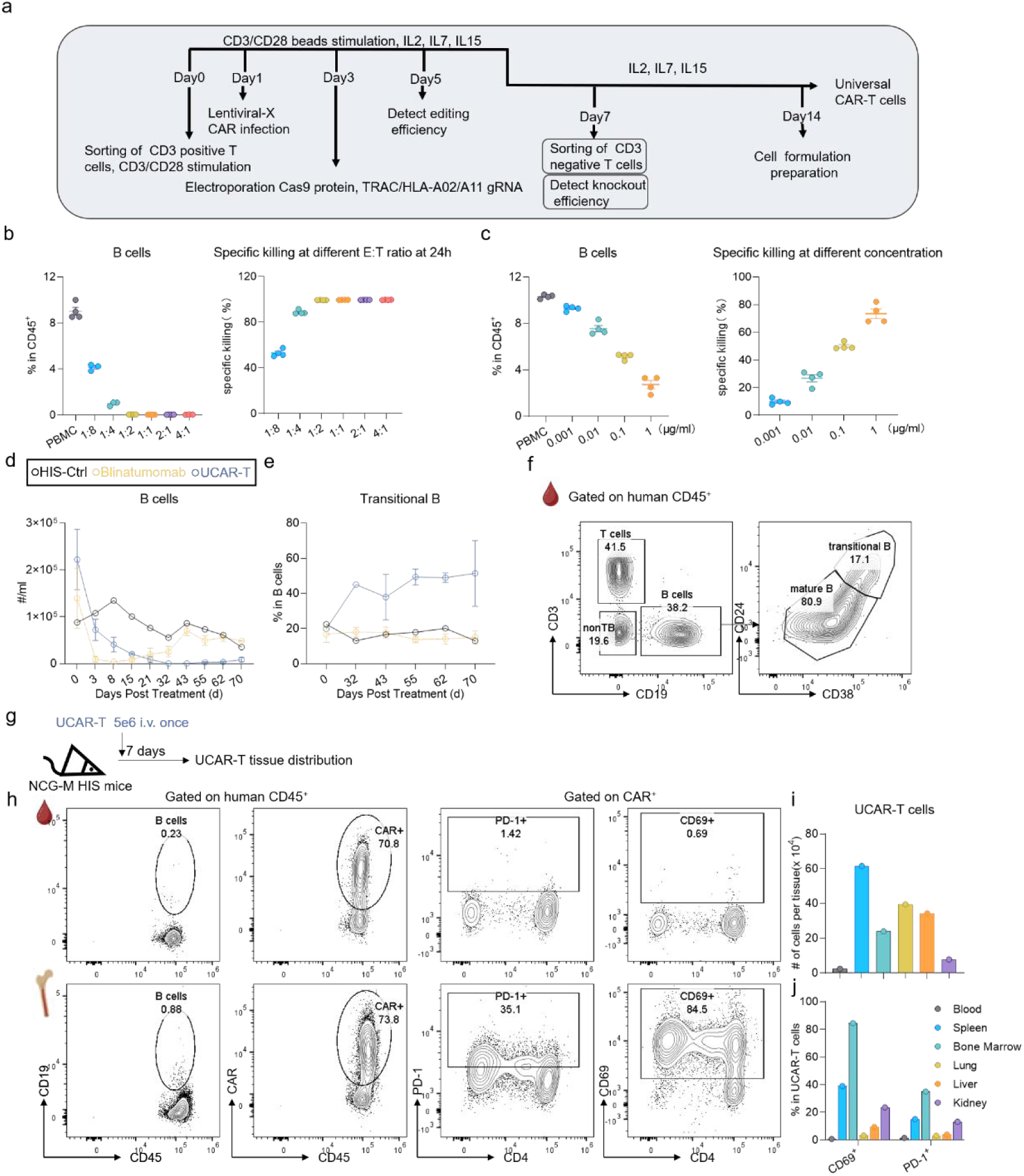
Quality control of allogeneic CAR-T cells. (a) The schematic diagram of UCAR-T cells preparation. (b)Allogeneic CAR-T cells were cocultured with PBMCs for 24 hours at indicated effector (UCAR-T cells) to target (B cells) ratio. Frequencies of B cells and calculated specific killing efficiency were summarized. (c) PBMCs were cultured in complete medium for 24 hours with different concentrations of CD3-CD19 bispecific antibody. Frequencies of B cells and calculated specific killing efficiency were summarized. (d) Flowcytometry analysis of absolute numbers of B cells in peripheral blood at indicated time points. (e-f) Representative plots(e) and summarized data(f) of frequencies of transitional B cells (CD24^hi^CD38^hi^) in peripheral blood at indicated time points (related to Figure 5E). (g) The schematic diagram of allogeneic CAR-T cells transfer and analysis. Five million UCAR-T cells were transferred to HIS-NCG-M mice and tissue infiltrated cells were analyzed one week later. (h-j) Phenotypic analysis of UCAR-T cells in different tissues. (h)Representative plots showing B cells depletion, UCAR-T cells infiltration and the activation (CD69^+^) and exhaustion (PD-1^+^) state of UCAR-T cells in peripheral blood(up) or bone marrow tissues(down). (i-j) Absolute numbers of UCAR-T cells(i) and frequencies of CD69^+^ and PD-1^+^ cells(j) were summarized. Data show mean ± SEM.

**Figure S7(related to Figure 5):**
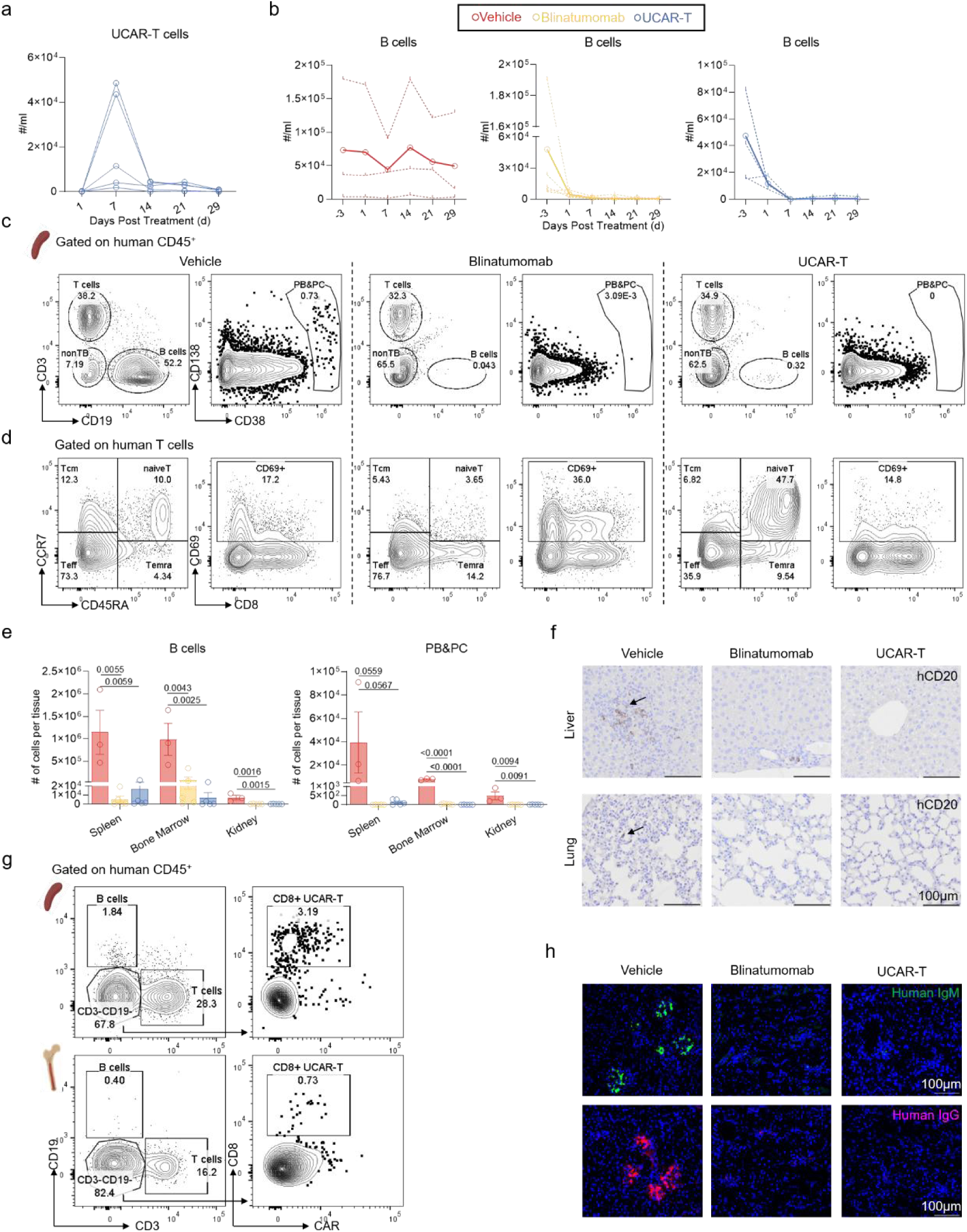
(a) Flowcytometry analysis of numbers of UCAR-T cells in peripheral blood at indicated time points. (b) Flowcytometry analysis of numbers of B cells in peripheral blood from three groups. Solid lines depicted the average level of each group and dotted lines showed the level of each individual. (c) Representative plots showing the gating of B cells, plasmablasts and plasma cells (PB&PC) in spleen tissues from Vehicle (left), Blinatumomab (mid) and UCAR-T (right) group. (d) Representative plots showing the frequencies of four T cell subsets and activated T cells in spleen tissues from Vehicle (left), Blinatumomab (mid) and UCAR-T (right) group (related to Figure 5L&5M). (e) Summarized data of Figure S7c. (f) Immunohistochemistry staining of human CD20 in the liver(up) and lung(down) tissues from three groups. Scale bar, 100μm. (g) Representative plots of residual UCAR-T cells in the spleen(up) and bone marrow(down) tissues one month after transfer. (h) Immunofluorescence staining of human IgM (up) and human IgG (down) deposition in the glomerular area of kidneys from three groups. Scare bar, 100μm. Data show mean ± SEM. Statistical analysis was performed using one-way ANOVA with Tukey multiple-comparison test. The exact *p* values are shown.

**Supplemental Table 1.** Clinical characteristics of SLE patients and healthy donors.

|  | **SLE(n=8)** | **HC(n=8)** |
| --- | --- | --- |
| **Age** | 35.38±8.05 (24~51) | 35.38±9.32 (26~56) |
| **Sex (% female)** | 7(87.50%) | 7(87.50%) |
| **WBC(×10<sup>9</sup>/L)** | 3.80±2.12 (2.3~7.4) |  |
| **RBC (×10<sup>12</sup>/L)** | 3.46±0.70 (2.31~4.57) |  |
| **Hb (g/L)** | 105.50±19.78 (73~138) |  |
| **PLT (×10<sup>9</sup>/L)** | 99.50±44.55 (68~291) |  |
| **C3 (g/L)** | 0.56±0.32 (0.2~1.2) |  |
| **C4 (g/L)** | 0.13±0.12 (0.02~0.39) |  |
| **Anti-dsDNA (IU/mL)** | 378.84±226.59<br>(147.17~856.095) |  |
| **ANA<sup>+</sup> (%)** | 8(100%) |  |
| **Anti-Sm<sup>+</sup> (%)** | 3(37.50%) |  |
| **Anti-U1RNP<sup>+</sup> (%)** | 4(50.00%) |  |
| **Urine protein(mg/L)** | 1973.28±1047.61<br>(487.1~3198.6) |  |

**Supplemental Table 2.** Antibodies used for FACS analysis.

| Antigen | Reactivity | Clone | Supplier | Dilution (50µl) |
| --- | --- | --- | --- | --- |
| CD45 | Human | HI30 | eBioscience | 1 µL |
| CD45 | Mouse | 30-F11 | eBioscience | 0.5 µL |
| CD3 | Human | UCHT1 | Biolegend | 1 µL |
| CD19 | Human | HIB19 | BD Bioscience | 1 µL |
| CD24 | Human | ML5 | Biolegend | 1 µL |
| CD38 | Human | HIT2 | Biolegend | 1 µL |
| CD45RA | Human | HI100 | BD Bioscience | 1 µL |
| CCR7 | Human | 3D12 | BD Bioscience | 1 µL |
| IFN-γ | Human | 4S.B3 | Biolegend | 1 µL |
| IL-17A | Human | BL168 | Biolegend | 1 µL |
| IL-21 | Human | 3A3-N3 | Biolegend | 1 µL |
| HLA-DR | Human | L243 | Biolegend | 1 µL |
| CD10 | Human | HI10a | Biolegend | 1 µL |
| CD27 | Human | L128 | BD Bioscience | 1 µL |
| IgD | Human | IA6-2 | Biolegend | 1 µL |
| IgG | Human | G18-145 | BD Bioscience | 1 µL |
| CD69 | Human | FN50 | Biolegend | 1 µL |
| CD123 | Human | 6H6 | eBioscience | 1 µL |
| CD138 | Human | MI15 | Biolegend | 1 µL |
| CD4 | Human | SK3 | BD Bioscience | 1 µL |
| CD8 | Human | RPA-T8 | Biolegend | 1 µL |
| CD20 | Human | 2H7 | Biolegend | 1 µL |
| TCRαβ | Human | IP26 | eBioscience | 1 µL |
| HLA-A2 | Human | BB7.2 | BD Bioscience | 1 µL |
| PD-1 | Human | EH12.2H7 | Biolegend | 1 µL |
| FMC63 | Human | Y45 | Acro<br>Biosystems | 0.5 µL |
| Eflour 780 |  |  | eBioscience | 0.05 µL |
| Eflour 506 |  |  | eBioscience | 0.125 µL |

